# A versatile toolkit for molecular QTL mapping and meta-analysis at scale

**DOI:** 10.1101/2020.12.18.423490

**Authors:** Corbin Quick, Li Guan, Zilin Li, Xihao Li, Rounak Dey, Yaowu Liu, Laura Scott, Xihong Lin

## Abstract

Molecular QTLs (xQTLs) are widely studied to identify functional variation and possible mechanisms underlying genetic associations with diseases. Larger xQTL sample sizes are critical to help identify causal variants, improve predictive models, and increase power to detect rare associations. This will require scalable and accurate methods for analysis of tens of thousands of molecular traits in large cohorts, and/or from summary statistics in meta-analysis, both of which are currently lacking. We developed APEX (All-in-one Package for Efficient Xqtl analysis), an efficient toolkit for xQTL mapping and meta-analysis that provides (a) highly optimized linear mixed models to account for relatedness and shared variation across molecular traits; (b) rapid factor analysis to infer latent technical and biological variables from molecular trait data; (c) fast and accurate trait-level omnibus tests that incorporate prior functional weights to increase statistical power; and (d) compact summary data files for flexible and accurate joint analysis of multiple variants (e.g., joint/conditional regression or Bayesian finemapping) without individual-level data in meta-analysis. We applied the methods to data from three LCL eQTL studies and the UK Biobank. APEX is open source: https://corbinq.github.io/apex.

## Introduction

Human genetics studies have identified tens of thousands of molecular QTLs-genetic loci associated with differences in molecular quantitative traits-including mRNA (eQTL), microRNA (miQTL), or protein (pQTL) expression, metabolite (metQTL), methylation (mQTL) levels (*1, 2*). By mapping DNA sequence variation to heritable differences in the transcriptome and epigenome, xQTL studies have provided important insights into genome function and gene regulation (*3–5*). xQTLs are also of interest in genome-wide association studies (GWAS) as possible biological antecedents of genetic associations with complex traits and diseases (*6–10*). Integrative analyses of xQTL and GWAS data have provided insight into the biological mechanisms underlying GWAS associations, and helped identify causal disease genes and therapeutic targets (*11–13*).

Larger xQTL studies are crucial to identify causal variants driving xQTL association signals, detect low-frequency and rare xQTL variants, and more accurately predict expression or methylation levels from genotype data. The next generation of xQTL studies will require scalable methods for association analysis in large multi-ethnic cohorts, accurate methods for downstream statistical analysis (e.g., Bayesian finemapping and colocalization analysis) from summary statistics in meta-analysis, and integrative methods to utilize prior knowledge of genome function. We developed APEX, a toolkit for scalable xQTL association analysis and meta-analysis, to address these challenges.

Molecular trait data suffers from a high degree of technical and biological variation, which can both mask and confound *cis* and *trans* genetic associations (*14–17*). Latent variable models such as PEER (*16*) and dimension reduction techniques such as principal component analysis (PCA) (*18, 19*) are often used to infer unobserved common sources of technical and biological variation in xQTL studies. PEER is particularly effective in xQTL analysis, but computationally demanding. In APEX, we implemented simple, efficient algorithms for high-dimensional factor analysis using early stopping for regularization (*20, 21*). We found that this approach is nearly as fast as PCA and far faster than PEER, while yielding equal or greater numbers of *cis* discoveries than either method.

Linear mixed models (LMM) are widely used to account for population structure and cryptic familial relatedness in genome-wide association studies (GWAS), and can additionally account for shared technical and biological variation across molecular traits in xQTL studies (*18*). However, despite multiple existing LMM methods for xQTL analysis (*18, 22*), ordinary least squares (OLS) is often used in practice for its greater computational efficiency. Even family-based eQTL studies often use a two-stage approach in which LMM residuals are used as response variables in OLS (*23, 24*), which may reduce statistical power (*25*). In APEX, we developed efficient algorithms for LMM association analysis to account for population structure, relatedness, and technical variation with tens of thousands of traits, which are accurate for small samples and scale linearly in sample size.

Permutation tests are the current standard to calculate trait-level xQTL omnibus tests and account for correlations between tests statistics across variants and traits in xQTL discovery (*19, 26, 27*). This approach is burdensome for large sample sizes, and does not readily capitalize prior knowledge of variant functionality. The aggregated Cauchy association test (ACAT) is a recently-developed method to combine p-values under arbitrary dependence structures (*28, 29*). We applied ACAT to aggregate xQTL test statistics for each molecular trait, which scales linearly in the number of variants and independent of sample size. Unlike permutation tests, which implicitly assign equal prior weight to all variants, ACAT can incorporate functional prior weights between variants and molecular traits. We found that simply weighting by the chromosomal distance between each variant and transcription start site (TSS) (*30*) substantially increased xQTL discoveries.

While dozens of xQTL studies have been conducted (*2*), meta-analysis is hampered by difficulties sharing human genomic data. Marginal variant-trait associations can be meta-analyzed using regression slopes and standard errors or z-scores alone. However, these statistics are not sufficient for analyses that involve the joint effects of multiple variants, such as joint and conditional analysis (*31, 32*), Bayesian finemapping (*33–37*), aggregation tests (*31, 38, 39*), and colocalization analysis (*40*). Multiple-variant analysis further requires variance-covariance or linkage disequilibrium (LD) matrices, which characterize the joint distribution of single-variant xQTL association statistics. In GWAS, proxy LD from a genotype reference panel is often used for multiple-variant analysis from summary statistics, but this is problematic for small or ancestrally heterogenous samples (*32, 35*), both of which are common in omics studies (*2, 3, 17, 41*). Indeed, previous xQTL meta-analyses have generally analyzed only marginal variant-trait associations (*42–44*). In APEX, we developed compact xQTL summary association data formats for accurate multiple-variant analysis in meta-analysis without individual-level data.

## Results

### Software development

We developed APEX (All-in-one Package for Efficient Xqtl analysis), a software toolkit for scalable xQTL mapping and meta-analysis. Core running modes for molecular trait preprocessing, *cis* and *trans* association analysis, and xQTL meta-analysis are summarized in Figure 1 (see Methods and Supplementary Materials for further details). APEX is a command-line tool implemented in C++, supports multithreading to expedite linear algebra, and provides flexible sub-setting options to facilitate parallelization across genomic regions. It uses the Eigen (*45*) and Spectra (*46*) C++ libraries for linear algebra, and HTSlib to process indexed BED, BCF, and VCF files (*47*). Precompiled Linux binaries and source code are available online.

**Figure 1.**
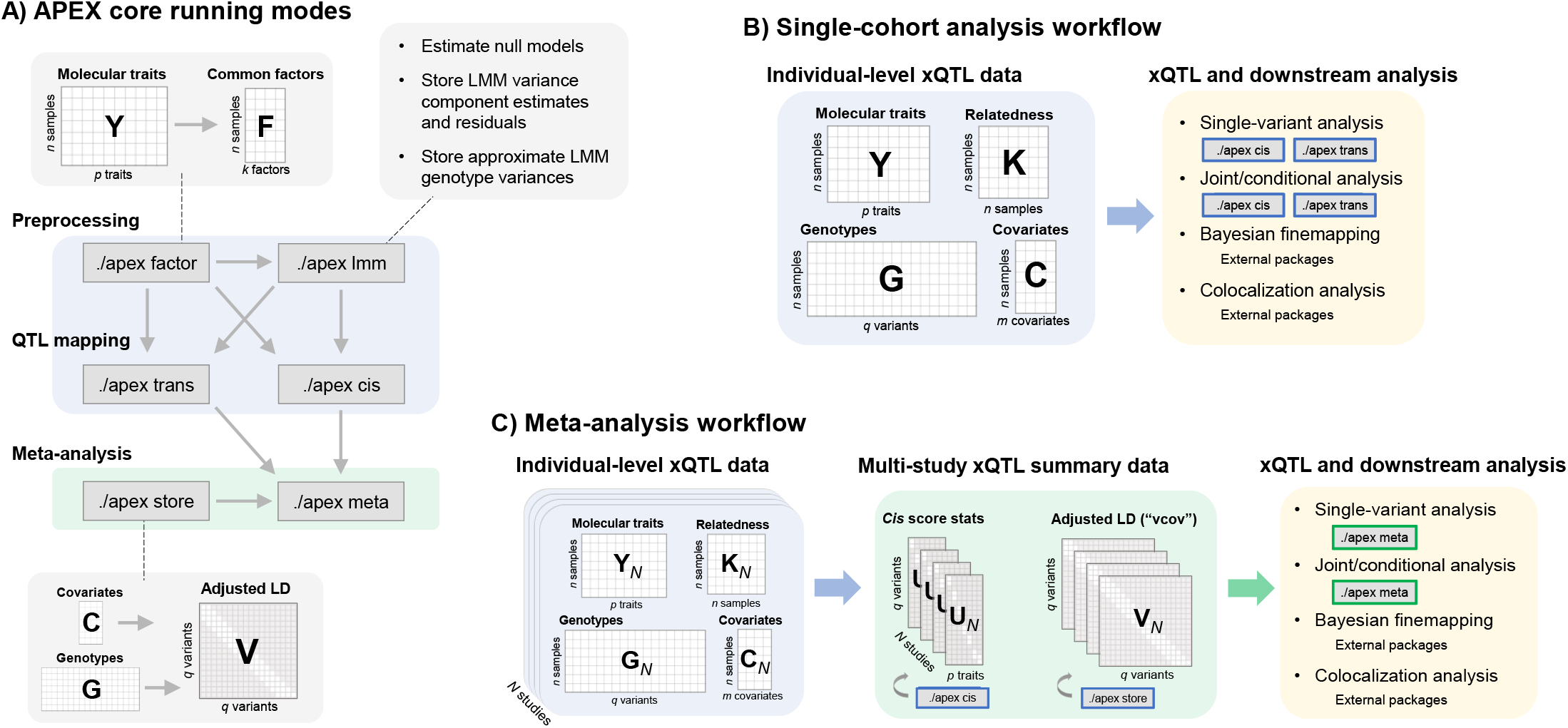
APEX toolkit for molecular QTL mapping and meta-analysis. **A:** Mode *factor* provides factor analysis to infer shared technical and biological factors across traits. In QTL mapping (modes *cis* and *trans*), inferred factor covariates can be modeled as fixed effects (by appending matrix **F** to covariate matrix **C**) or random effects (using mode *lmm*). Mode *lmm* enables rapid linear mixed model (LMM) association analysis (in modes *cis* and *trans*) by precomputing and storing variance component estimates, LMM trait residuals, and approximate LMM genotypic variances. Mode *store* generates compact adjusted LD files for accurate multiple-variant analysis from summary statistics (using mode *meta* for meta-analysis). **B:** Individual-level molecular trait, genotype, and covariate data (and optional genetic relatedness matrix) are used as input for single-variant and joint/conditional association analysis across traits (APEX modes *cis* and *trans*). These data can also be used for Bayesian finemapping and colocalization analysis using external software packages. **C:** Each study generates summary data files (single-variant score statistics using mode *cis* and adjusted LD matrices using mode *store*) from individual-level data. These summary files can be used for single-variant and joint/conditional association meta-analysis in mode *meta*, or combined using the *Apex2R* interface to create input data for Bayesian finemapping and colocalization analysis using external packages.

### Application to 3 lymphoblastoid cell line (LCL) eQTL data sets

We analyzed LCL eQTLs using genotype, expression, and technical covariate data from the GTEx project v8 (*41*), Geuvadis project (*5*), and HapMap project (*3, 48, 49*) (Table 1). GTEx (n = 147) and Geuvadis (n = 454) have RNA-seq LCL expression measurements and whole genome sequencing (WGS) based genotype calls. HapMap (n = 518) has array-based LCL expression measurements and array-based genotype calls, from which we imputed genotypes using the 1000 Genomes Project reference panel (*50*). Data and processing procedures for each study are further described in Methods.

**Table 1.**
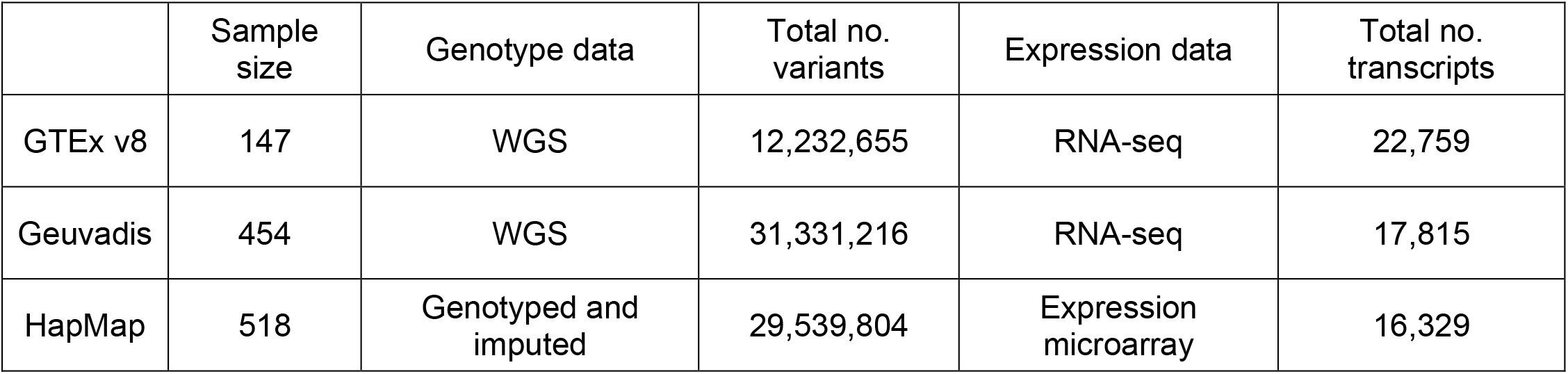
Descriptive statistics for LCL eQTL data sets. Summary of LCL data sets analyzed. For HapMap, we report the number of imputed variants. For all studies, we report the number of variants before filtering. Processing and filtering procedures for each study are described in Methods.

### Rapid factor analysis of molecular traits for xQTL analysis

We inferred hidden covariates from gene expression measurements in each study using PEER (*16*), expression principal component (ePC) analysis (*19*), and expression factor analysis (eFA) (*20, 21*). For each method, we varied the number of hidden covariates from 1 to 100. eFA and PEER explicitly model shared and unique variances for each trait, whereas ePCs capture maximal variance across all traits (*51*). Conceptually, ePC can be viewed as a special case of eFA in which all traits are assumed to have equal unique variance (unexplained by common factors). Further details are given in Methods and Supplementary Materials.

We used APEX to perform *cis*-eQTL analysis in each study modeling the hidden factor covariates as either fixed effects using ordinary least squares (OLS) or random effects using restricted maximum likelihood (REML) (*14*) (Figure 2). ePC and eFA covariates were calculated directly in APEX, and PEER factors were calculated using the PEER R package (*16*). For each method and data set, we varied the number of inferred covariates between 1 and 100. Across the studies, APEX eFA was 86 to 5033 times faster than PEER for models with >50 common factors (and 30 to 779 times faster for 20 to 50 factors), and provided equal or greater numbers of *cis* discoveries in each of the 3 data sets (Figure 2, panel A). Random-effect eFA provided the greatest number of discoveries in each of the 3 data sets, and fixed-effect or random-effect ePCs generally yielded the smallest numbers of discoveries.

**Figure 2.**
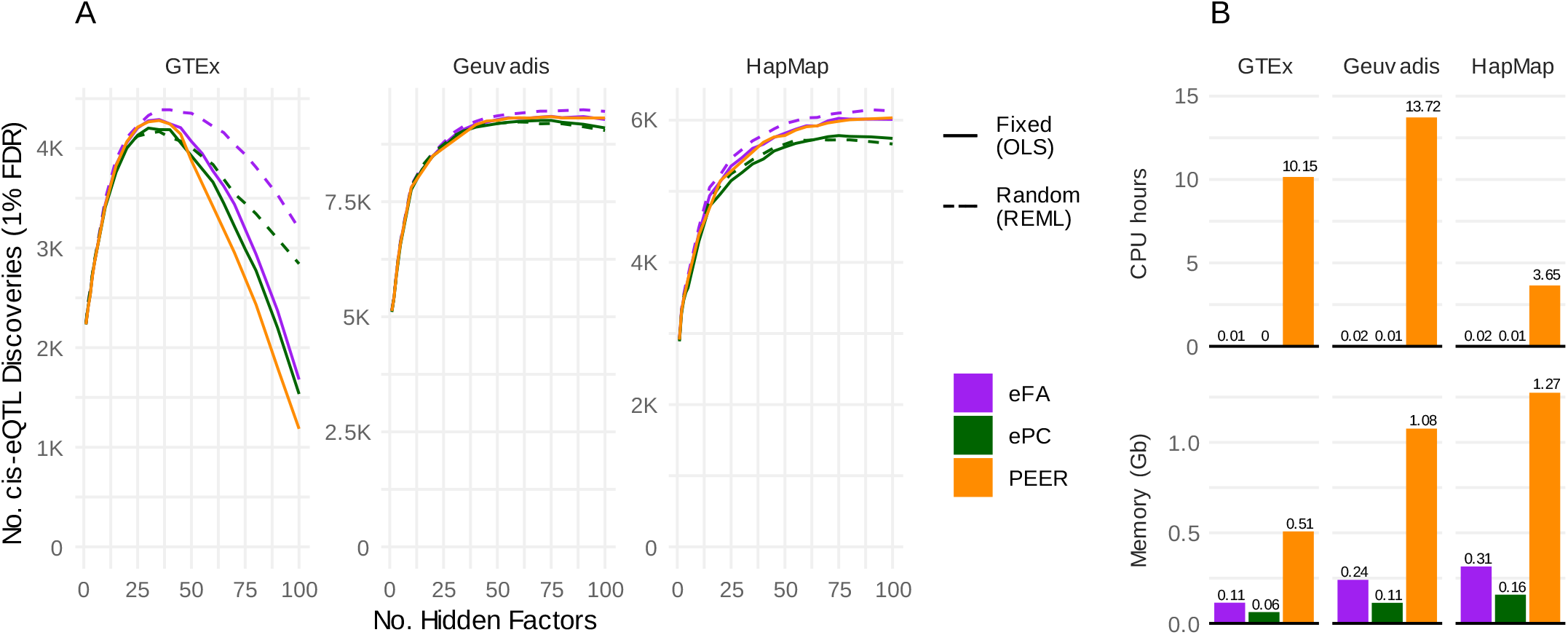
Rapid factor analysis and linear mixed models for *cis*-eQTL analysis. **A:** Number of LCL cis-eQTL discoveries at 1% FDR as a function of the number of hidden factors (x axis) inferred using PEER, factor analysis (eFA), or principal components analysis (ePC) across 3 studies. ePC and eFA covariate effects were estimated either as fixed effects (using OLS) or random effects (using REML) in association analysis using APEX. PEER covariates effects were estimated as fixed effects. **B:** Total running time (CPU hours) and maximum memory usage to generate ePC, eFA, and PEER covariates across models with 5, 10, 20, 40, 60, 80, and 100 latent factors. All jobs used a single CPU core. ePC and eFA covariates were calculated using APEX; PEER covariates were calculated using the PEER R package version 1.3 with a maximum of 1000 iterations.

To assess Type I error rates for fixed-effect and random-effect models with ePC or eFA covariates, we simulated 100 expression data sets under the null hypothesis in the Geuvadis study. We used the empirical covariance between expression and observed covariates (not inferred from expression) and empirical variance matrix of expression residuals (projecting out observed covariates) to simulate expression under the null hypothesis matching the observed covariance structure (Supplementary Figures 1-2). With each simulated expression matrix, we re-calculated the inferred covariates (eFA or ePC) and performed *cis*-eQTL analysis modeling the inferred covariates as either fixed or random effects. Association tests from all configurations (fixed-effect or random-effect models with between 1 and 100 inferred covariates) showed well-calibrated Type I error rates (Supplementary Figure 3).

### Fast, scalable linear mixed models with tens of thousands of molecular traits

We assessed the computational performance and numerical concordance of APEX and standard tools for linear mixed model (LMM) association analysis: FastGWA (*52*), BOLT-LMM (*53*), GMMAT (*54*), and GENESIS (*55*). APEX uses a 3-stage approach to efficiently estimate LMM null models and association statistics with tens of thousands of traits (Supplementary Figure 4), whereas the other tools are intended for single-trait analysis. We note that each of these tools supports a variety of features not considered in our analysis here—for example, GMMAT and GENESIS support flexible generalized LMM (GLMM) for binary and other non-normal traits, and BOLT-LMM supports flexible variance partitioning. For LMM association analysis, FastGWA and BOLT-LMM use approximations for efficient analysis in large cohorts, which may be less accurate with smaller sample sizes (e.g., < 5000 (*56*)). GENESIS, GMMAT, and APEX do not use such approximations, and APEX further uses small-sample LMM association tests (Supplementary Materials). To assess computational performance for LMM association analysis in large cohorts, we used genotype data and a sparse GRM for 10,000 individuals from the UK Biobank study, and simulated expression data for 16,329 traits with heritability drawn from a uniform distribution (Methods). Variant component estimates and single-variant association test statistics were nearly numerically equivalent between APEX, GMMAT, and GENESIS, as expected; FastGWA test statistics showed lower concordance with other methods (Supplementary Figure 5). LMM association analysis using APEX was >200-fold faster than GENESIS and GMMAT, 51.4-fold faster than BOLT-LMM, and 2.5-fold faster than FastGWA (Supplementary Table 1).

### Powerful and efficient *cis*-xQTL omnibus tests

We performed single-variant and gene-level *cis*-eQTL analysis in each study using APEX, FastQTL, and QTLtools (Figure 3). APEX and FastQTL use multiple linear regression (MLR) to adjust for covariates by default, whereas QTLtools uses simple linear regression with expression residuals (SLR-resid). We note that QTLtools can also perform MLR by regressing out covariates from genotype files prior to association analysis. Gene-level p-values from QTLtools and FastQTL use a Beta-approximated permutation test (Beta), whereas APEX uses either ACAT with constant weights (ACAT) or ACAT with distance-to-TSS weights between each variant and gene (ACAT-dTSS). FastQTL was run using adaptive p-values with 100 to 1000 permutations; QTLtools was run with 1000 permutations.

**Figure 3.**
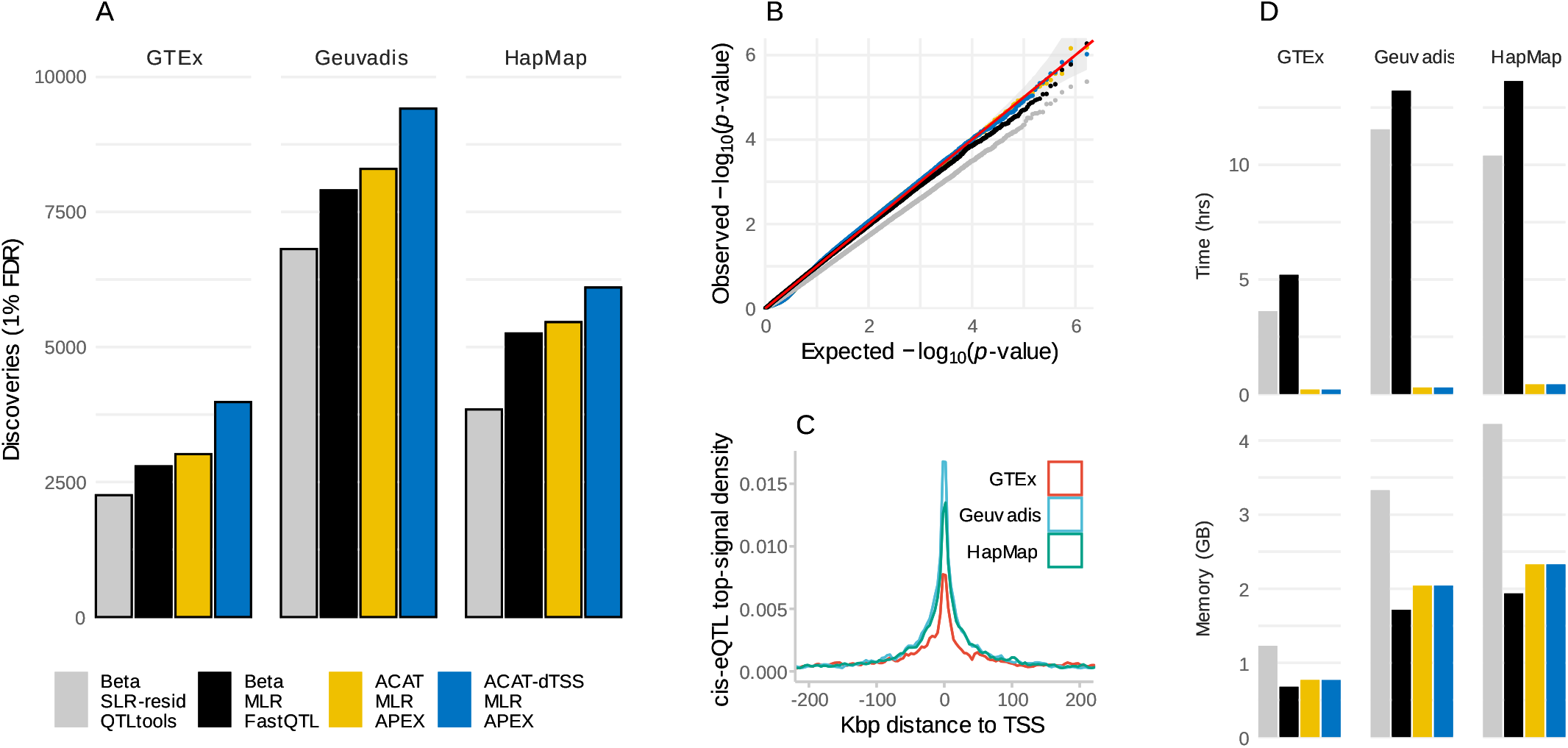
Fast and powerful *cis*-eQTL omnibus tests. **A**: **ACAT and dTSS weights increase eGene discoveries.** Gene-level *cis*-eQTL discoveries for each LCL data set at 1% FDR. Because all methods maintain calibrated Type I error rates in simulations (panel B), a larger number of discoveries suggests greater statistical power. Note that the number of tested genes varies across the three studies (Figure 4). **B**: **Calibration of permutation-based and ACAT p-values.** Q-Q plots for each method in simulations under the null hypothesis using genotype and expression data from Geuvadis. Traits were simulated using the observed correlation structure of gene expression, and expression PC covariates were re-calculated from simulated expression values in each replicate (Methods). P-values for all methods maintain calibrated or conservative Type I error rates, and SLR-resid permutation-based p-values are notably conservative. **C: eQTL enrichment by dTSS.** Density of chromosomal distance between top cis-eVariant and TSS across genes for each study. Cis-eVariants are strongly enriched nearer the TSS. **D: CPU time and memory for eGene discovery.** Analyses were run sequentially across chromosomes with 1 CPU; we report maximum memory usage and total elapsed running time.

We compared the numbers of *cis*-eQTL discoveries at 1% false discover rate (FDR) in each study from Beta permutation tests using FastQTL (*27*) or QTLtools (*19*), and from ACAT (*28, 29*) using APEX (Figure 3 panel A). Each method calculates gene-level omnibus *cis*-eQTL p-values (*cis*-eGene p-values) based on single-variant association test statistics within a 1 megabase (Mbp) window of the transcription start site (TSS). QTLtools and FastQTL use permutation tests of the minimum p-value across variants, and expedite computation by modeling the null distribution as a beta density using a fixed number of permutations (*27*). In each of the three studies, ACAT and permutation-based p-values were generally concordant (Supplementary Figure 6), but ACAT yielded more *cis*-eGene discoveries overall and was >30x faster (Figure 3, panels A and D). We also calculated weighted ACAT test statistics, in which each variant received a weight proportional to *e^−γ|d|^* where *d* is the number of base pairs between the variant and TSS and *γ* = 1e-5 (*30*). dTSS weighting further increased the number of *cis*-eGene discoveries by 14 to 30% across single studies (Figure 3, panel A).

We assessed p-value calibration for ACAT (implemented in APEX) and permutation-based p-values (implemented in FastQTL and QTLtools) by simulating expression data under the null hypothesis using genotype and expression data from the Geuvadis study (Figure 3 panel B). We used the sample covariance matrices of expression and observed covariates to simulate expression traits under the observed covariance structure (Methods). Empirical Type I error rates were well-controlled for both ACAT and Beta p-values, and SLR-resid p-values were conservative (shown previously in (*57*)). Permutation test p-values from SLR-resid were also notably conservative, which is expected because while trait residuals and genotype residuals are orthogonal to covariates, permuted trait residuals and unadjusted genotypes are not.

### Accurate multiple-variant xQTL meta-analysis from summary statistics

We assessed CPU time, memory, and storage required to create summary files for xQTL meta-analysis using APEX. We generated single-variant association summary statistics (sumstat files) and adjusted LD matrices (vcov files, which store the variance-covariance of association test statistics) for each of the 3 studies using APEX (Supplementary Figures 7-8). Summary statistics files were generated across all autosomes in 0.17 to 0.33 CPU hours and required 0.42 to 0.49 Gb storage per study (Supplementary Table 2). Adjusted LD files, which included LD for all pairs of variants within sliding 2 Mbp windows, were generated across all autosomes in 32.1 to 75.3 CPU hours and required 21.5, 34.3, 119.7 GB storage for GTEx, Geuvadis, and HapMap respectively (Supplementary Table 2). HapMap, which used imputed genotype dosages, required notably more time and storage than the other studies, which used WGS-based hard-call genotypes. We also compared adjusted LD storage using RareMetalWorker (RMW) (*31*), a tool for rare-variant association meta-analysis, across the 3 studies. APEX was 1.5 to 2.2-fold faster and required 4.5 to 21.5-fold less storage than RMW (Supplementary Table 3).

Score statistics and adjusted LD (stored in APEX sumstat and vcov files) are sufficient for a wide range of analyses involving the joint effects of multiple variants, including joint and conditional analysis, Bayesian finemapping, and penalized linear regression. We used APEX sumstat and vcov files from each LCL study to perform stepwise regression analysis using APEX-meta (Figure 4 and Supplementary Figure 9) and Bayesian finemapping using the *susieR* R package (*33*) (Figure 5) in individual studies and meta-analysis. To assess the accuracy of summary-based analyses, we also performed these analyses from individual-level data. Stepwise regression slopes and p-values and finemapping posterior inclusion probabilities (PIPs) were nearly numerically equivalent between individual-level vs sumstat data (Pearson Rsq > 0.999; Figure 5 panel C).

**Figure 4.**
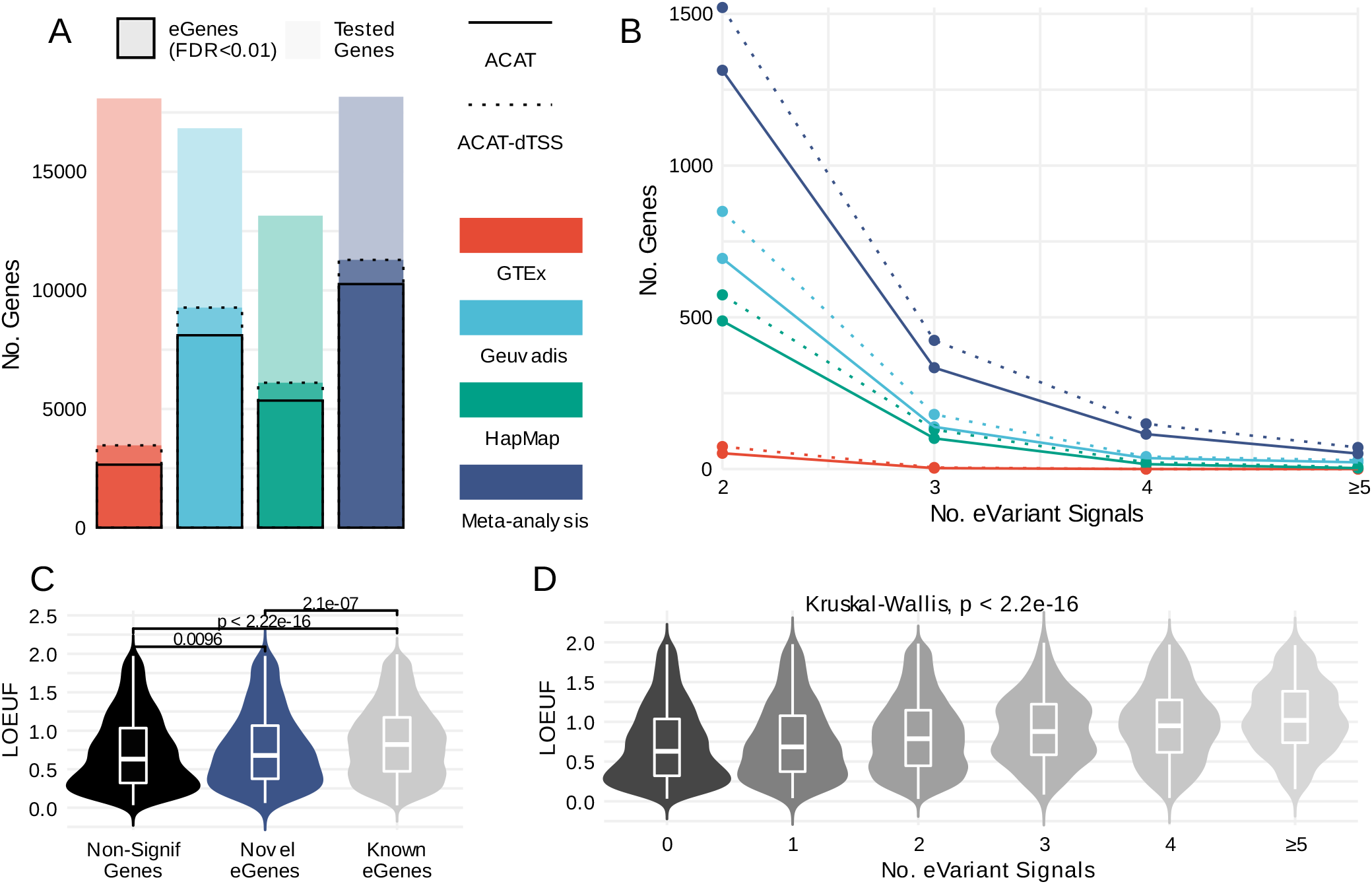
Meta-analysis identifies novel primary and secondary *cis*-eQTLs. **A: Meta-analysis and dTSS weights increase eGene discoveries.** eGenes detected in LCL cis-eQTL analysis across studies and meta-analysis. Colored bars show total numbers of tested genes, and outlined bars show numbers of eGenes (cis-eQTL genes) detected at 1% FDR using unweighted ACAT (solid line) and or distance to transcription start site (dTSS) weighted ACAT (dashed line). dTSS weights increased eGene discoveries by 30.6% for GTEx, 14.4% for Geuvadis, 14.1% for HapMap, and 10.0% for meta-analysis. **B: Meta-analysis and dTSS weights increase secondary eQTL discoveries.** Secondary cis-eQTL variant discoveries across studies and meta-analysis. Shown are numbers of genes with 2, 3, 4, or ≥5 LCL eQTL eVariant signals detected at 1% FDR using unweighted (solid line) and dTSS-weighted ACAT. dTSS weights increased secondary signal discoveries by 43.6% for GTEx, 23.3% for Geuvadis, 20.4% for HapMap, and 19.3% for meta-analysis. **C: Meta-analysis detects** *cis***-eQTLs for constrained genes**. Loss of function (LoF) observed/expected upper bound fraction (LOEUF) is a metric of genetic constraint; constrained genes have smaller LOEUF. LOEUF densities are shown for the 11,750 genes present in all (3 out of 3) studies, divided into 3 categories: (a) no cis-eQTLs detected at 1% FDR (2,659 “non-signif” genes), (b) ≥1 eQTL detected in meta-analysis but not individual studies (693 “novel eGenes”), and (c) ≥1 eQTL detected by ≥1 individual study (8,398 “known eGenes”). Both novel and non-significant genes have significantly lower LOEUF than known eGenes, suggesting greater constraint. **D: Fewer secondary** *cis***-eQTLs are detected for constrained genes.** LOEUF densities for genes with 0, 1, … ≥5 significant eVariants detected by stepwise regression in meta-analysis (1% FDR), shown for genes present in 3 out of 3 studies. Genes with more eVariants tend to have higher LOEUF (less constraint), as expected.

**Figure 5.**
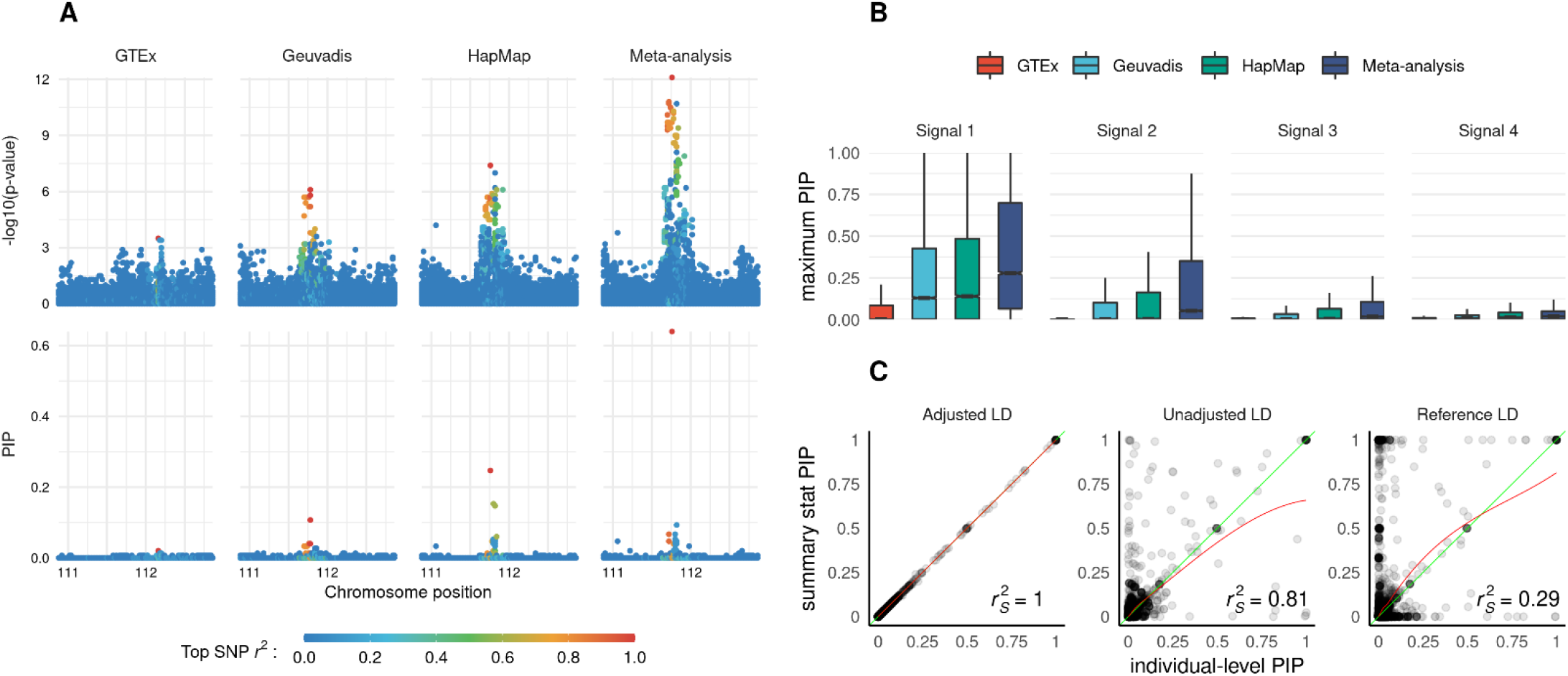
Accurate QTL finemapping from summary statistics. APEX xQTL sumstat and vcov files enable accurate multiple-variant analyses without individual-level data. Here, we illustrate Bayesian finemapping from APEX summary statistics data using the *susieR* package and *Apex2R* interface to access sumstat and vcov files. **A: Finemapping** *cis***-eQTLs from summary statistics.** *cis*-eQTL p-values (upper panel) and posterior inclusion probabilities (PIPs) for *cis* variants at the *FYN* locus (6.p22) are shown across the three studies and meta-analysis. Meta-analysis increases signal strength (upper panels) and precision identifying putative causal variants (lower panels). **B: Meta-analysis increases finemapping precision.** We finemapped 9,787 genes present each of the 3 studies from APEX sumstat and vcov summary data files using the *susieR* package. For each gene, we assigned each variant to its most likely signal cluster (highest posterior probability), and calculated the maximum PIP across variants within each signal cluster. Boxplots show the distribution of the maximum PIP within the 1st, 2nd, 3rd and 4th signal cluster across genes for each study. Maximum PIPs tend to increase with sample size, as expected. **C: APEX sumstat and vcov files enable accurate finemapping from summary statistics.** Concordance of PIPs across 71 genes using individual-level data (x axis) vs summary statistics (y axis) from HapMap with covariate-adjusted HapMap LD (left), HapMap LD not adjusted for covariates (middle), or proxy LD from Geuvadis (right) adjusted for similar covariates. PIPs from summary statistics using APEX vcov files (adjusted LD) are nearly numerically equivalent with individual-level analysis. PIPs using unadjusted or proxy LD are less concordant with individual-level analysis (Spearman *r*^2^ 0.81 or 0.29 respectively).

To assess the accuracy of joint analysis from association summary statistics using proxy LD or unadjusted LD rather than APEX vcov files (which store adjusted LD), we performed finemapping with association summary statistics from HapMap and either (a) unadjusted LD (the correlation matrix of genotypes in HapMap, not adjusted for PCs and other covariates), or (b) proxy LD (adjusted LD from Geuvadis as a proxy for adjusted LD from HapMap). Unadjusted LD is often used for multiple-variant analysis from GWAS summary statistics (e.g., (*32*)), and differs from adjusted LD when genotypes are correlated with covariates (e.g., genotype PCs in multi-ethnic studies). PIPs using proxy LD or unadjusted LD yielded substantially lower concordance with the exact PIPs that adjusted LD (Figure 5 panel C), which is expected due to the relatively small sample sizes and differences in ancestry composition between HapMap and Geuvadis. Notably, many other xQTL studies have relatively small sample size and heterogeneous ancestry composition (Supplementary Figure 10).

### Functional characterization of LCL eQTL variants and genes

We hypothesized that mRNA expression heritability is lower for genes that are more evolutionarily constrained, and that therefore eGenes detected only in meta-analysis are more constrained on average than those detected in single studies. To assess this hypothesis, we compared the loss-of-function observed/expected upper bound fraction (LOEUF), a recently developed metric of genetic constraint (smaller LOEUF suggests greater constraint) (*58*), across genes that were tested in all 3 studies (11,750 genes). Novel LCL eGenes (eQTL associations detected by meta-analysis, but not by individual studies) and genes with no significant signal had significantly lower LOEUF than previously-identified eGenes (Mann–Whitney p = 2.1e-7 and 2.2e-16 respectively), while the difference in LOEUF was less pronounced for novel eGenes vs genes with no detected eQTLs (p = 0.0096) (Figure 4 panel C). Moreover, genes with larger numbers of significant *cis*-eQTL signals (identified in stepwise regression; Methods) tend to have larger LOEUF values (p < 2.2e-16) (Figure 4 panel D). While gene length is associated with LOEUF, we observed no significant trends between gene length and eQTL signals. These results suggest that larger samples sizes will be required to detect xQTLs for more biologically important genes, highlighting the utility of meta-analysis.

We assessed functional enrichment of primary and secondary LCL eQTL variants identified in meta-analysis across the 3 studies. We used binomial logistic regression to identify features associated with LCL eQTL variants controlling for distance to nearest TSS and minor allele frequency (MAF) (Methods). First, we assessed enrichment of LCL eQTL variants in tissue-specific DNase I hypersensitive sites (DHSs) across 16 tissue groups (*59*). LCL eQTLs showed striking enrichment in lymphoid-specific DHS compared to other tissue groups (Supplementary Figure 11). Next, we assessed overlap enrichment of LCL eQTL variants overlapping GWAS variants identified using the NHGRI-EBI GWAS Catalog (*60*). Among 15 categories of GWAS traits, LCL eQTL variants showed strongest enrichment with GWAS variants for immune diseases (Supplementary Figure 12). These results suggest that LCL eQTL variants capture cell-type specific functionality, and highlight the utility of xQTL analysis in diverse tissues and cell types.

## Discussion

Future xQTL studies will be conducted in increasingly large and diverse cohorts, and are poised to capitalize on growing knowledge of functional elements in the human genome. We developed APEX to empower these studies by providing a flexible, scalable framework for *cis* and *trans* xQTL analysis and meta-analysis. APEX provides rapid high-dimensional factor analysis to infer latent technical and biological factors, efficient linear mixed model (LMM) association analysis for *cis* and *trans* xQTL mapping, procedures to incorporate prior weights in primary and secondary xQTL signal discovery, and a framework for accurate joint analysis of multiple variant effects from xQTL summary data.

Our LMM framework for molecular traits extends upon previous work (*14, 22*) by optimizing association analysis with high-dimensional traits and structured random-effect covariance matrices. In particular, we precompute and recycle computationally expensive terms for each molecular trait and each variant, and exploit the structure of random-effect covariance matrices (low-rank or block-diagonal) to expedite linear algebra. With these optimizations, LMM association analysis scales linearly in sample size and the number of traits, enabling rapid analysis with large xQTL cohorts. APEX also uses small-sample adjustment and avoids large-sample approximations to provide accurate p-values for smaller cohorts.

In GWAS, random effects are typically used to account for infinitesimal genetic effects or familial relatedness in LMM association analysis. In xQTL studies, random effects can also be used to model shared technical and biological variation across traits (*14, 22*). Our results suggest that this strategy outperforms ordinary least squares (OLS) when using expression factor analysis covariates, but underperforms OLS when using expression PC covariates. A variety of other methods can be applied to infer hidden covariates from molecular trait data, and various other strategies (e.g., penalized regression) can be used to include these covariates in xQTL analysis. We believe this is a worthy area for further research. Here, our work provides rapid inference of latent technical and biological covariates from molecular trait data, and a flexible LMM framework to include these covariates as fixed or random effects in xQTL association analysis.

Our meta-analysis framework extends from previous eQTL meta-analysis tools (*61*) by enabling accurate multiple-variant analysis, including joint/conditional analysis (using APEX mode meta), Bayesian finemapping (using *susieR* (*33*) or DAP (*34*)), and colocalization analysis (using external software), from xQTL summary statistics. These methods are fundamental in a variety of applications, including predictive weight estimation (e.g., for TWAS) and integrative analysis of GWAS loci. Methods that use LD from a reference panel as a proxy for meta-analysis LD may be inaccurate when reference or meta-analysis sample size is limited (e.g., < 5000), ancestry composition differs between reference vs meta-analysis samples, or genotypes are correlated with covariates in meta-analysis. In APEX, we provide exact study-specific adjusted LD matrices (vcov files); similar strategies have been used for rare-variant association meta-analysis (*31, 38*), but not to our knowledge for genome-wide xQTL or finemapping meta-analysis. The proposed xQTL meta-analysis framework enables flexible and highly accurate multiple-variant modeling with arbitrary sample sizes, ancestry compositions, and sets of covariates.

While our applications focused on eQTL studies, APEX sumstat and vcov formats are also well-suited for GWAS of quantitative traits, which can be used, for example, in colocalization analysis of GWAS and xQTL signals. More broadly, we encourage GWAS and xQTL studies to publicly release adjusted LD data in addition to single-variant association summary statistics when possible. With streamlined tools for the analysis of such data, greater availability of sufficient statistics including LD would increase reproducibility, enhance meta-analysis, and accelerate discovery.

The statistical methods in APEX can be extended in a variety of ways, such as by (a) leveraging correlations between molecular traits across multiple tissues or cell-types, (b) modeling genetic correlations between traits of the same tissue or cell-type, or (c) supporting generalized linear models for non-normal traits. Multivariate LMMs can be used to account for the correlation structure of genetic and environmental components of molecular traits across and within tissues or cell-types. Also, zero-inflated Poisson or negative binomial generalized linear mixed models (GLMMs) may be desirable for some types of molecular trait data.

Our data applications have several limitations, including (a) analysis of only LCL eQTLs, (b) relatively small eQTL sample sizes, and (c) limited *trans*-eQTL analysis. Our LCL eQTL analysis revealed striking enrichment with relevant tissue-specific DHS, highlighting the utility of xQTL analysis across diverse tissues and cell types. Moreover, APEX is well suited for analysis of mRNA expression and other molecular traits across broader sets of tissues or cell types due to its computational efficiency. While the three LCL eQTL had limited sample sizes, our simulation studies using UK Biobank genotype data demonstrated that APEX is scalable to larger cohorts, with >100-fold improvement in CPU time relative to standard tools. Finally, we note that APEX fully supports *trans*-eQTL analysis, as illustrated in simulation studies.

In summary, APEX provides an efficient and comprehensive framework for *cis* and *trans* xQTL mapping and meta-analysis. For xQTL studies of a single cohort, APEX provides efficient inference of latent technical and biological factors from molecular trait data (*20*), which performs competitively with state-of-the-art methods in *cis*-eQTL analysis and orders of magnitude faster; rapid LMM association analysis with tens of thousands of molecular traits; powerful, efficient trait-level xQTL omnibus tests; and accurate multiple-variant analysis. For xQTL meta-analysis, APEX provides accurate single-variant and joint multiple-variant regression analysis, and compact summary data formats for flexible and accurate multiple-variant modeling (e.g., Bayesian finemapping) without individual-level data across multiple studies.

## Online Methods

### Statistical methods implemented in APEX

#### Principal components and factor analysis of molecular traits

APEX provides efficient algorithms to calculate principal components (PCA) and factor analysis (FA) of molecular traits. For PCA, we calculate *k* PC covariates as the first *k* left singular vectors of the truncated singular value decomposition (SVD) of the *n* × *p* normalized expression matrix **Y**, which is scaled and centered so that each column (trait) has mean 0 and variance 1. The SVD is **Y** = **UDV**^T^, and **U**_(*k*)_ = (*U*_1_, *U*_2_, …, *U*_*k*_) are the PC covariates. When the number of traits is larger than the number of samples, we calculate **U**_(*k*)_ from the truncated SVD (or eigendecomposition) of **YY**^T^, as **YY**^T^ = **UD**^2^**U**^T^. Otherwise, we calculate 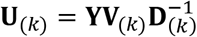, where the right singular vectors **V**_(*k*)_ are calculated from **Y**^T^**Y** = **VD**^2^**V**^T^.

The FA model is **Y** = **ZB** + **E** where **Z** is the *n* × *k* matrix of common factors, **B** is the *k* × *p* matrix of factor loadings, and **E** is the *n* × *p* matrix of unique factors. The rows of **E** are independent, and each row vector is multivariate normal with covariance matrix 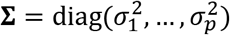. In APEX, we estimate the common factors **Z** using an SVD of 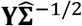, which we initialize with constant variances 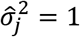 for all *j* = 1, 2, …, *p*. Given the first *k* left singular vectors 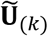 of 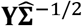, we update the estimates as 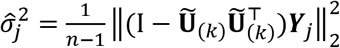 for each trait *j* = 1, 2, …, *p*, and repeat. A similar algorithm was suggested by (*62*), but the underlying likelihood is unbounded if 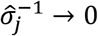, and convergence generally fails in practice. As proposed by (*20, 21*), we perform regularization by halting after a fixed number of iterations. If the number of samples is greater than the number of traits (*n* > *p*), we modify this approach using the *p* × *k* right singular vectors rather than the *n* × *k* left singular vectors of 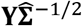 The time complexity of this procedure is *O*(min(*n*, *p*)^2^ *k* + *pnk*), where *n* is the sample size, *p* is the number of traits, and *k* is the number of factors. Further details are given in Supplementary Materials.

#### Statistical methods for cis and trans LMM association analysis

APEX provides a scalable linear mixed model (LMM) framework to account for familial relatedness (*14, 63*) or technical variation (*18, 22*) (Supplementary Figure 4). For traits *t* = 1, 2, …, *p*, we assume the model

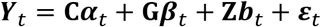

where **Y**_*t*_ is the observed trait, **C** is the matrix of fixed-effect covariates, **G** is the matrix of genotypes, and **Z** is the matrix of random-effect covariates. To account for relatedness, **ZZ**^T^ = **K** where **K** is a genetic relatedness matrix (GRM); and to account for technical and biological variation, **Z** is comprised of inferred factor covariates. We assume the residual *ε*_*t*_ is multivariate normal distributed with mean **0** and variance 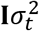, and the random effects are multivariate normally distributed with mean **0** and variance 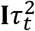.

By default, variance components are estimated by restricted maximum likelihood (REML) under the null hypothesis of no single-variant associations. APEX supports sparse (*64, 65*) and low-rank (*66*) covariance matrices for random effects, and uses specialized optimizations for each structure. We expedite computation by precomputing and saving variance component estimates and LMM residuals for each trait, and residual genotypic variance terms for each variant. While APEX precomputes LMM residuals, we note that it does not use the GRAMMAR-gamma (*67*) or related approximations. For *trans*-xQTL analysis (considering all variant-trait pairs), the time complexity of LMM estimation and association testing in APEX is *O*(*pm*^2^*n* + *npq* + *nmq*) where *n* is the sample size, *p* the number of traits, *m* the number of covariates, and *q* the number of variants. Further details are provided in Supplementary Materials.

#### Omnibus p-values for cis-xQTL signals

We used the aggregated Cauchy association test (ACAT) (*28, 29*) to calculate omnibus *cis* region p-values for primary and secondary signals. ACAT omnibus p-values are calculated as *p*^*O*^ = *F*{∑_*i*_ *w*_*i*_*F*^−1^(*p*_*i*_)} where *F* is the cumulative density function (CDF) of the standard Cauchy distribution, *w*_*i*_ are non-negative weights with ∑_*i*_ *w*_*i*_ = 1, and *p*_*i*_ are p-values. ACAT provides valid p-values under arbitrary dependence structures, provided that *p*_*i*_ are valid p-values (calibrated under the null hypothesis). When *p*_*i*_ are single-variant p-values in the *cis* region, we find that ACAT p-values with constant weights are highly concordant with permutation-based p-values (Supplementary Figure 6), but much faster (Figure 3, Panel B).

#### Data formats for flexible and accurate xQTL meta-analysis

APEX provides genetic association summary statistics (sumstat) and variance-covariance (vcov) data in an indexed, compressed binary format (Supplementary Figures 7-8). For fixed effects models, APEX sumstat files store the vector of score statistics *U*_*t*_ = **G**^T^**PY**_**t**_ and residual sum of squares **Y**_*t*_^T^ **PY**_*t*_ for each trait *t*, where **G** is the genotype matrix, **P** is a projection matrix, and **Y** is the matrix of molecular traits; APEX vcov files store the variance-covariance matrix of score statistics **V** = **G**^T^**PG** (also called adjusted LD matrix). For *cis* analysis, we store only score statistics for variants within a window of each molecular trait (1 Mbp by default), and adjusted LD for variants within twice the specified window size. These statistics are sufficient for a wide variety of downstream statistical analyses (for example, multiple-variant joint and conditional regression modeling, aggregation tests, Bayesian finemapping, and colocalization analysis), and preserve the genetic privacy of xQTL study participants. Similar strategies have been used to aggregate variants for gene-based tests in rare-variant (RV) GWAS meta-analysis (*31, 38*), but to our knowledge no existing methods exist for efficiently sharing and combining adjusted LD for genome-wide meta-analysis of common variants in GWAS or xQTL studies. APEX summary data can be combined across multiple studies for meta-analysis in APEX mode *meta* for joint and conditional regression analysis, or accessed and combined through an R interface for use with other packages. Further details are given in Supplementary Materials.

#### Secondary xQTL signal discovery

We implemented stepwise regression algorithms to detect multiple conditionally independent genetic association signals (Supplementary Figure 9) using either individual-level data or sumstat and vcov files. At each iteration, we evaluate signal-level significance using an omnibus p-value to test the null hypothesis that no remaining variants are associated with the trait, calculated as *p*^*O*^ = *F*{∑_*j*∈*U*_ *w*_*j*_*F*^−1^(*p*_*j*|*S*_)}, where *S* and *U* are the current sets of selected and unselected variants, *p*_*j*|*S*_ is the conditional p-value for variant *j* given selected variants *S*, *w*_*j*_ is the weight for variant *j* (normalized so that ∑_*j*∈*U*_ *w*_*j*_ = 1 at iteration), and *F* is the CDF of the standard Cauchy distribution. If *p*^*O*^ < *α*, where *α* is a specified threshold, we select the most significant variant in *U* (adding it to *S* and removing it from *U*) and continue; otherwise, we retain the current set *S* and exit. Further details and extensions are given in Supplementary Materials.

### Data sources

#### LCL eQTL genotype data

Genotype data from the 1000 Genomes Project Phase 3 in NCBI build 38 were obtained from the International Genome Sample Resource (IGSR) webpage (*68*). WGS-based genotype data for the GTEx project v8 were obtained from dbGaP under accession number (phg 001219.v1); variants and samples with >15% missingness were excluded. Remaining missing genotype calls were imputed as best-guess hard call genotypes using the phasing software Eagle (*69*). Genotype data from the HapMap project in NCBI build 36 from the Broad Institute webpage. This data set included 1,379,607 autosomal variants; to increase the number of variants overlapping the other studies, HapMap genotypes were imputed with the 1000 Genomes Project Phase 3 reference panel using Minimac3 (*70*); imputed variants were filtered with Mach-Rsq > 0.3. A final list of 10,930,386 variants, the intersection of variants across the three studies, was used for meta-analysis. Kinship matrices and genetic principal component covariates were calculated using PLINK 2 (*64*).

#### LCL gene expression data

RNA-seq expression data from the Geuvadis consortium, which performed RNA-seq on LCLs for a subset of samples in the 1000 Genomes Project, were obtained from the IGSR webpage (*5*). RNA-seq expression data from LCLs for GTEx v8 participants were obtained from dbGaP under accession number (phe000037.v1). LCL expression microarray data for 618 individuals in the HapMap 3 study (*17*) were obtained from ArrayExpress (*71*); Illumina probe identifiers were mapped to Ensembl gene identifiers using the illuminaHumanv2 Bioconductor R package. Genes that were lowly expressed (count ≤ 5) in ≥ 25% of individuals were excluded. Expression microarray measurements and RNA-seq TPMs were rank-normal transformed within each study (*5*).

#### UK Biobank genotype data

Genotype data from the UK Biobank study were obtained under Application Number 52008. UK Biobank protocols were approved by the National Research Ethics Service Committee and written informed consent were signed by the participants. Marker variants were filtered by including only autosomal SNPs with genotype missingness < 1% that passed all batch-wise genotype quality control steps (*72*) (590,606 variants after filtering). We randomly selected a multi-ethnic subset of 10,000 UK Biobank participants for analysis, among which 4,000 were Irish, 3,000 were South Asian (Indian, Pakistani, and Bangladeshi), and 3,000 were African and Caribbean (all self-reported). We generated an ancestry-adjusted sparse genetic relatedness matrix (GRM) using LD-pruned MAF > 0.01 variants in R (*73*) by projecting out genotype PCs from genotypes and setting GRM elements to 0 for >4th degree estimated relatives (genetic correlation < 0.044). LD pruning used pairwise *r*^2^ < 0.1 in sliding windows of 50 SNPs moving 5 SNPs at a time.

### Data analysis and simulation procedures

#### Molecular trait simulation procedures

To evaluate Type I error rates of association test statistics, we simulated expression data under the null hypothesis of no single-variant genetic associations in the Geuvadis study. We used the empirical covariance between expression and technical covariates and simulate covariance of expression residuals to simulate expression with a realistic correlation structure (Supplementary Figures 1-2). Specifically, in each replicate, we simulated the row vector of expression across genes for participant *i* as a multivariate normal distribution with mean 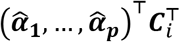 and variance 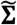, where **C**_*i*_ is the *i*^*th*^ row vector of from technical covariates **C** (genotype PCs, gender, batch, ethnicity indicator), 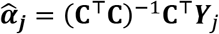 is the estimated effects of technical covariates on gene *j* expression **Y**_*j*_ (column vector), and 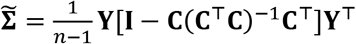 is the sample covariance matrix of expression residuals across genes. In each simulation replicate, we re-calculated the inferred covariates (ePC, eFA, or PEER) from the simulated expression matrix.

We simulated expression data in the UK Biobank study to assess the computational performance of linear mixed models (LMMs) for xQTL analysis in large cohorts, which will be critical to identify rare and small-effect xQTL variants and molecular traits that contribute to heritable diseases. In these experiments, we simulated each trait independently from a multivariate normal distribution with mean **Cα**, where **C** is the matrix of genotype PCs, and variance *h*^2^**K** + (1 − *h*^2^)**I** where **K** is the sparse genetic relatedness matrix. We simulated the covariate effects **α** from an independent normal distribution, and pseudo-heritability parameter *h*^2^ from a uniform distribution.

#### LCL eQTL enrichment analysis

We used binomial logistic regression models to assess functional enrichment of LCL eQTLs. The mean model was specified 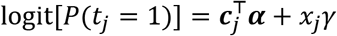, where the outcome was defined as *t*_*j*_ = 1 if variant *j* is in high LD (*r*^2^ > 0.8) with a lead LCL eQTL variant for any gene and *t*_*j*_ = 0 otherwise, where lead eQTL variants were identified using stepwise regression (described above). The scalar *x*_*j*_ denotes the feature of interest (e.g., *x*_*j*_ = 1 if variant *j* overlaps a lymphoid-specific DHS and *x*_*j*_ = 0 otherwise), and the covariate vector **c**_*j*_ included an intercept and cubic b-spline terms for log-transformed minor allele frequency (MAF) and distance to nearest transcription start site (TSS). We included all variants that were tested for *cis* association (within 1 Mbp of TSS for any tested gene).

## Contributions

Conception: CQ

Conceptualization: CQ, LG, LS, X Lin

Primary software development: CQ, LG

Software and data formats: CQ, LG, ZL, X Li

Statistical methods: CQ, LG, RD, YL, X Lin

Data acquisition and preparation: CQ, LG, RD, X Lin

LCL eQTL analysis: CQ, LG

UK Biobank analysis: CQ, RD

Simulation studies: CQ

Figures: CQ

Primary manuscript writing: CQ, LG

Manuscript editing and review: All authors

## Acknowledgements

We thank Hyun Min Kang for helpful discussions on linear mixed models and factor analysis. We thank Andy Shi for assistance with UK Biobank data, and Sheila Gaynor for helpful discussions on GTEx data.

The Genotype-Tissue Expression (GTEx) Project was supported by the Common Fund of the Office of the Director of the National Institutes of Health, and by NCI, NHGRI, NHLBI, NIDA, NIMH, and NINDS. The data used for the analyses described in this manuscript were obtained from dbGaP accession numbers no. phe000037.v1 and phg 001219.v1 on 11/22/2019.

This research has been conducted using the UK Biobank Resource under Application Number 52008.

## References

1. B. Ng et al., An xQTL map integrates the genetic architecture of the human brain’s transcriptome and epigenome. Nature neuroscience 20, 1418 (2017).

2. Z. Zheng et al., QTLbase: an integrative resource for quantitative trait loci across multiple human molecular phenotypes. Nucleic acids research 48, D983–D991 (2020).

3. J. T. Bell et al., DNA methylation patterns associate with genetic and gene expression variation in HapMap cell lines. Genome biology 12, R10 (2011).

4. L. Parts, O. Stegle, J. Winn, R. Durbin, Joint genetic analysis of gene expression data with inferred cellular phenotypes. PLoS Genet 7, e1001276 (2011).

5. T. Lappalainen et al., Transcriptome and genome sequencing uncovers functional variation in humans. Nature 501, 506–511 (2013).

6. E. R. Gamazon et al., Predixcan: Trait mapping using human transcriptome regulation. BioRxiv, 020164 (2015).

7. P. Zeng, T. Wang, S. Huang, Cis-SNPs set testing and PrediXcan analysis for gene expression data using linear mixed models. Scientific reports 7, 1–11 (2017).

8. A. Gusev et al., Transcriptome-wide association study of schizophrenia and chromatin activity yields mechanistic disease insights. Nature genetics 50, 538–548 (2018).

9. N. Mancuso et al., Large-scale transcriptome-wide association study identifies new prostate cancer risk regions. Nature communications 9, 1–11 (2018).

10. Y. Zhang et al., PTWAS: investigating tissue-relevant causal molecular mechanisms of complex traits using probabilistic TWAS analysis. Genome biology 21, 1–26 (2020).

11. H. Fang et al., A genetics-led approach defines the drug target landscape of 30 immune-related traits. Nature genetics 51, 1082–1091 (2019).

12. I. Tachmazidou et al., Identification of new therapeutic targets for osteoarthritis through genome-wide analyses of UK Biobank data. Nature genetics 51, 230–236 (2019).

13. Á. Duffy et al., Tissue-specific genetic features inform prediction of drug side effects in clinical trials. Science Advances 6, eabb6242 (2020).

14. H. M. Kang et al., Variance component model to account for sample structure in genome-wide association studies. Nature genetics 42, 348–354 (2010).

15. O. Stegle, L. Parts, R. Durbin, J. Winn, A Bayesian framework to account for complex non-genetic factors in gene expression levels greatly increases power in eQTL studies. PLoS Comput Biol 6, e1000770 (2010).

16. O. Stegle, L. Parts, M. Piipari, J. Winn, R. Durbin, Using probabilistic estimation of expression residuals (PEER) to obtain increased power and interpretability of gene expression analyses. Nature protocols 7, 500 (2012).

17. B. E. Stranger et al., Patterns of cis regulatory variation in diverse human populations. PLoS Genet 8, e1002639 (2012).

18. H. M. Kang, C. Ye, E. Eskin, Accurate discovery of expression quantitative trait loci under confounding from spurious and genuine regulatory hotspots. Genetics 180, 1909–1925 (2008).

19. O. Delaneau et al., A complete tool set for molecular QTL discovery and analysis. Nature communications 8, 1–7 (2017).

20. A. B. Owen, J. Wang, Bi-cross-validation for factor analysis. Statistical Science 31, 119–139 (2016).

21. J. Wang, Ph. D. thesis, Stanford University, (2016).

22. A. Ziyatdinov et al., lme4qtl: linear mixed models with flexible covariance structure for genetic studies of related individuals. BMC bioinformatics 19, 1–5 (2018).

23. A. Buil et al., Gene-gene and gene-environment interactions detected by transcriptome sequence analysis in twins. Nature genetics 47, 88–91 (2015).

24. E. Grundberg et al., Mapping cis-and trans-regulatory effects across multiple tissues in twins. Nature genetics 44, 1084–1089 (2012).

25. J. Eu-Ahsunthornwattana et al., Comparison of methods to account for relatedness in genome-wide association studies with family-based data. PLoS Genet 10, e1004445 (2014).

26. G. Consortium, The GTEx Consortium atlas of genetic regulatory effects across human tissues. Science 369, 1318–1330 (2020).

27. H. Ongen, A. Buil, A. A. Brown, E. T. Dermitzakis, O. Delaneau, Fast and efficient QTL mapper for thousands of molecular phenotypes. Bioinformatics 32, 1479–1485 (2016).

28. Y. Liu et al., Acat: A fast and powerful p value combination method for rare-variant analysis in sequencing studies. The American Journal of Human Genetics 104, 410–421 (2019).

29. Y. Liu, J. Xie, Cauchy combination test: a powerful test with analytic p-value calculation under arbitrary dependency structures. Journal of the American Statistical Association 115, 393–402 (2020).

30. C. Quick, X. Wen, G. Abecasis, M. Boehnke, H. M. Kang, Integrating Comprehensive Functional Annotations to Boost Power and Accuracy in Gene-Based Association Analysis. BioRxiv, 732404 (2019).

31. S. Feng, D. Liu, X. Zhan, M. K. Wing, G. R. Abecasis, RAREMETAL: fast and powerful meta-analysis for rare variants. Bioinformatics 30, 2828–2829 (2014).

32. J. Yang et al., Conditional and joint multiple-SNP analysis of GWAS summary statistics identifies additional variants influencing complex traits. Nature genetics 44, 369–375 (2012).

33. G. Wang, A. Sarkar, P. Carbonetto, M. Stephens, A simple new approach to variable selection in regression, with application to genetic fine mapping. Journal of the Royal Statistical Society: Series B (Statistical Methodology), (2020).

34. Y. Lee, F. Luca, R. Pique-Regi, X. Wen, Bayesian Multi-SNP Genetic Association Analysis: Control of FDR and Use of Summary Statistics. bioRxiv, 316471 (2018).

35. C. Benner et al., Prospects of fine-mapping trait-associated genomic regions by using summary statistics from genome-wide association studies. The American Journal of Human Genetics 101, 539–551 (2017).

36. X. Wen, Y. Lee, F. Luca, R. Pique-Regi, Efficient integrative multi-SNP association analysis via deterministic approximation of posteriors. The American Journal of Human Genetics 98, 1114–1129 (2016).

37. C. Benner et al., FINEMAP: efficient variable selection using summary data from genome-wide association studies. Bioinformatics 32, 1493–1501 (2016).

38. X. Zhan, Y. Hu, B. Li, G. R. Abecasis, D. J. Liu, RVTESTS: an efficient and comprehensive tool for rare variant association analysis using sequence data. Bioinformatics 32, 1423–1426 (2016).

39. X. Li et al., Dynamic incorporation of multiple in silico functional annotations empowers rare variant association analysis of large whole-genome sequencing studies at scale. Nature genetics 52, 969–983 (2020).

40. X. Wen, R. Pique-Regi, F. Luca, Integrating molecular QTL data into genome-wide genetic association analysis: Probabilistic assessment of enrichment and colocalization. PLoS genetics 13, e1006646 (2017).

41. F. Aguet et al., The GTEx Consortium atlas of genetic regulatory effects across human tissues. BioRxiv, 787903 (2019).

42. Y. Kim et al., A meta-analysis of gene expression quantitative trait loci in brain. Translational psychiatry 4, e459–e459 (2014).

43. U. Võsa et al., Unraveling the polygenic architecture of complex traits using blood eQTL metaanalysis. BioRxiv, 447367 (2018).

44. S. K. Sieberts et al., Large eQTL meta-analysis reveals differing patterns between cerebral cortical and cerebellar brain regions. Scientific data 7, 1–11 (2020).

45. G. Guennebaud, B. Jacob, Eigen: a c++ linear algebra library. URL http://eigen.tuxfamily.org, Accessed 22, (2014).

46. Y. Qiu, Spectra C++ Library For Large Scale Eigenvalue Problems. URL https://spectralib.org/, (2020).

47. P. Danecek et al., The variant call format and VCFtools. Bioinformatics 27, 2156–2158 (2011).

48. R. A. Gibbs et al., The international HapMap project. (2003).

49. W. Zhang, M. J. Ratain, M. E. Dolan, The HapMap resource is providing new insights into ourselves and its application to pharmacogenomics. Bioinformatics and biology insights 2, BBI. S455 (2008).

50. G. P. Consortium, A global reference for human genetic variation. Nature 526, 68–74 (2015).

51. M. E. Tipping, C. M. Bishop, Probabilistic principal component analysis. Journal of the Royal Statistical Society: Series B (Statistical Methodology) 61, 611–622 (1999).

52. L. Jiang et al., “A resource-efficient tool for mixed model association analysis of large-scale data,” (Nature Publishing Group, 2019).

53. P.-R. Loh et al., Efficient Bayesian mixed-model analysis increases association power in large cohorts. Nature genetics 47, 284 (2015).

54. H. Chen, M. P. Conomos, GMMAT-package: Generalized Linear Mixed Model Association Tests. (2020).

55. S. M. Gogarten et al., Genetic association testing using the GENESIS R/Bioconductor package. Bioinformatics 35, 5346–5348 (2019).

56. P.-R. Loh, BOLT-LMM v2.3.4 User Manual. URL https://alkesgroup.broadinstitute.org/BOLT-LMM/downloads/BOLT-LMM_v2.3.4_manual.pdf (2019).

57. P. Yajnik, M. Boehnke, Power loss due to testing association between covariate‐adjusted traits and genetic variants. Genetic Epidemiology, (2020).

58. K. J. Karczewski et al., The mutational constraint spectrum quantified from variation in 141,456 humans. Nature 581, 434–443 (2020).

59. W. Meuleman et al., Index and biological spectrum of human DNase I hypersensitive sites. Nature, 1–8 (2020).

60. A. Buniello et al., The NHGRI-EBI GWAS Catalog of published genome-wide association studies, targeted arrays and summary statistics 2019. Nucleic acids research 47, D1005–D1012 (2019).

61. A. F. Di Narzo, H. Cheng, J. Lu, K. Hao, Meta-eQTL: a tool set for flexible eQTL meta-analysis. BMC bioinformatics 15, 392 (2014).

62. D. N. Lawley, Vi.—the estimation of factor loadings by the method of maximum likelihood. Proceedings of the Royal Society of Edinburgh 60, 64–82 (1940).

63. A. L. Price et al., Principal components analysis corrects for stratification in genome-wide association studies. Nature genetics 38, 904–909 (2006).

64. C. C. Chang et al., Second-generation PLINK: rising to the challenge of larger and richer datasets. Gigascience 4, s13742-13015-10047-13748 (2015).

65. A. Manichaikul et al., Robust relationship inference in genome-wide association studies. Bioinformatics 26, 2867–2873 (2010).

66. C. Lippert et al., FaST linear mixed models for genome-wide association studies. Nature methods 8, 833–835 (2011).

67. G. R. Svishcheva, T. I. Axenovich, N. M. Belonogova, C. M. van Duijn, Y. S. Aulchenko, Rapid variance components–based method for whole-genome association analysis. Nature genetics 44, 1166–1170 (2012).

68. S. Fairley, E. Lowy-Gallego, E. Perry, P. Flicek, The international genome sample resource (IGSR) collection of open human genomic variation resources. Nucleic Acids Research 48, D941–D947 (2020).

69. P.-R. Loh, P. F. Palamara, A. L. Price, Fast and accurate long-range phasing in a UK Biobank cohort. Nature genetics 48, 811–816 (2016).

70. S. Das et al., Next-generation genotype imputation service and methods. Nature genetics 48, 1284–1287 (2016).

71. A. Athar et al., ArrayExpress update–from bulk to single-cell expression data. Nucleic acids research 47, D711–D715 (2019).

72. C. Bycroft et al., The UK Biobank resource with deep phenotyping and genomic data. Nature 562, 203–209 (2018).

73. R. C. Team. (Vienna, Austria, 2013).

