## Supplementary Text for "A versatile toolkit for molecular QTL mapping and meta-analysis at scale"

### APEX Supplementary Materials

December 17, 2020

**References** **12**

#### 0.1 Key Formula and Notations

This supplement describes statistical procedures for analysis of quantitative molecular trait loci (xQTL) analysis implemented in APEX (All-in-one Package for Efficient Xqtl analysis). For molecular traits $t = 1, 2, ..., p$ across studies $s = 1, 2, ..., N$, we assume the model

$$\boldsymbol{Y}_{st} = \mathbf{C}_s \boldsymbol{\zeta}_{st} + \mathbf{G}_s \boldsymbol{\beta}_{st} + \mathbf{Z}_s \boldsymbol{b}_{st} + \boldsymbol{\varepsilon}_{st}, \tag{1}$$

where $\boldsymbol{Y}_{st}$ is the molecular trait, $\mathbf{C}_s$ is the matrix of technical covariates, $\mathbf{G}_s$ is the matrix of genotypes, $\mathbf{Z}_s$ is the matrix of random-effect covariates with coefficients $\boldsymbol{b}_{st} \sim \mathcal{N}(\mathbf{0}, \mathbf{I}\tau^2_{st})$, and $\boldsymbol{\varepsilon}_{st} \sim \mathcal{N}(\mathbf{0}, \mathbf{I}\sigma^2_{st})$ is the residual. We define the matrix $\boldsymbol{\Omega}_{st} = \text{Var}(\boldsymbol{Y}_{st}) = \mathbf{K}_s\tau^2_{st} + \mathbf{I}\sigma^2_{st}$ where $\mathbf{K}_s = \mathbf{Z}_s\mathbf{Z}_s^\top$. The random effects can be specified to account for familial relatedness across samples (when $\mathbf{K}_s$ is a genetic relatedness matrix), or to capture shared technical or biological variation across traits (Section 0.2). The subscripts $s$ and $t$ are omitted in later sections that discuss a single trait or study.

APEX provides single- and multiple-variant xQTL association analysis methods, as well as single-study ($N = 1$) and meta-analysis ($N > 1$), all of which are described by the above model. Single-variant xQTL association analysis, also called xQTL mapping, involves fitting models in which only a single element of $\boldsymbol{\beta}_s$ is non-zero (Section 0.4). Joint and conditional analysis involve fitting models in which a specified set of two or more elements of $\boldsymbol{\beta}_s$ are non-zero (Sections 0.5 and 0.6). Meta-analysis typically assumes $\boldsymbol{\beta}_s = \boldsymbol{\beta}_{s'}$ across all studies $s$ and $s'$, but can also allow for heterogenous genetic effects across studies (Section 0.6).

While LMM methods in APEX are scalable for large sample sizes (Sections 0.3 and 0.4.2), LMM association tests are also designed to be accurate for both small and large sample sizes (Section 0.4.1), as well as low-frequency and rare variants (Section 0.4.2). APEX also provides multiple-variant analysis methods to detect secondary QTL signals, and compact data formats for QTL association summary statistics and variance-covariance data to enable accurate, efficient multiple-variant analysis without individual-level data (Section 0.6).

#### 0.2 Rapid Factor Analysis of Molecular Traits

Let $\mathbf{Y} \in \mathbb{R}^{n\times p}$ denote the matrix of $p$ molecular traits measured across individuals $i = 1, 2, ...., n$. We consider the model $\mathbf{Y} = \mathbf{ZB} + \mathbf{E}$ where $\mathbf{Z} \in \mathbb{R}^{n\times k}$ are common factors, $\mathbf{B} \in \mathbb{R}^{k\times p}$ are factor loadings, and $\mathbf{E}$ are unique factors. We assume the rows of $\mathbf{Y}$, given by $\boldsymbol{Y}_i = \boldsymbol{Z}_i\mathbf{B} + \boldsymbol{E}_i$, are conditionally independent given $\mathbf{Z}$, and $\boldsymbol{E}_i^\top \sim \mathcal{N}(\mathbf{0}, \boldsymbol{\Sigma})$ where $\boldsymbol{\Sigma} = \text{diag}(\sigma_1^2, ..., \sigma_p^2)$. Then, $\mathbf{Y}$ follows the $n \times p$ matrix normal distribution with expectation $\mathbf{ZB}$, $n \times n$ row scale matrix $\mathbf{I}$, and $p \times p$ column scale matrix $\boldsymbol{\Sigma}$. The complete-data log likelihood (omitting constants) given $\mathbf{M} = \mathbf{ZB}$ is

$$\log \mathcal{L}(\boldsymbol{\Sigma}, \mathbf{M}) = -\frac{n}{2}\log|\boldsymbol{\Sigma}| - \frac{1}{2}\text{tr}\left\{\boldsymbol{\Sigma}^{-1}(\mathbf{Y} - \mathbf{M})^\top(\mathbf{Y} - \mathbf{M})\right\}, \tag{2}$$

$$\text{tr}\left\{\boldsymbol{\Sigma}^{-1}(\mathbf{Y} - \mathbf{M})^\top(\mathbf{Y} - \mathbf{M})\right\} = \sum_{i=1}^{n}\sum_{j=1}^{n}\sigma_j^{-2}(Y_{ij} - M_{ij})^2. \tag{3}$$

The maximum likelihood estimate of $\boldsymbol{\Sigma}$ given fixed $\mathbf{M}$ is $\hat{\sigma}_j^2 = \frac{1}{n}\sum_{i=1}^{n}(Y_{ij} - M_{ij})^2$ for $j = 1, 2, ..., p$, where $M_{ij} = \boldsymbol{z}_i^\top\boldsymbol{b}_j$. Also, the matrix $\mathbf{M}$ that maximizes $\log\mathcal{L}(\boldsymbol{\Sigma}, \mathbf{M})$ given fixed $\boldsymbol{\Sigma}$ and subject to the constraint rank$(\mathbf{M}) = k$ is $\hat{\mathbf{M}}_{(k)} = \mathbf{U}_{(k)}\mathbf{U}_{(k)}^\top\mathbf{Y}$, where $\mathbf{U}_{(k)}$ are the first $k$ left singular vectors of $\mathbf{Y}\boldsymbol{\Sigma}^{-1/2}$ (which follows from the matrix approximation lemma, because the weight matrix $\boldsymbol{\Sigma}$ is diagonal). Iterating between these two steps, as has been described by (Lawley, 1940) and others, generally fails, because $\mathcal{L}(\boldsymbol{\Sigma}, \mathbf{M}) \to \infty$ as any $\hat{\sigma}_j^2 \to 0$. Recently, (Wang, 2016; Owen and Wang, 2016) suggested to terminate the algorithm after a fixed number of iterations as a form of regularization. We use this approach in APEX, which we find performs competitively with more sophisticated factor analysis methods such as PEER (Stegle et al., 2012).

We note that an alternate expression of $\hat{\mathbf{M}}_{(k)}$, which is given in (Wang, 2016; Owen and Wang, 2016), is $\hat{\mathbf{M}}_{(k)} = \mathbf{U}_{(k)}\mathbf{D}_{(k)}\mathbf{V}_{(k)}^\top\boldsymbol{\Sigma}^{1/2}$ where $\mathbf{U}_{(k)}\mathbf{D}_{(k)}\mathbf{V}_{(k)}^\top$ is the rank-$k$ truncated SVD of $\mathbf{Y}\boldsymbol{\Sigma}^{-1/2}$. The expressions are equivalent because $\mathbf{U}_{(k)}^\top\mathbf{Y} = \mathbf{D}_{(k)}\mathbf{V}_{(k)}^\top\boldsymbol{\Sigma}^{1/2}$. To show this, denote the full SVD by $\mathbf{Y}\boldsymbol{\Sigma}^{-1/2} = \mathbf{UDV}^\top$, partition the matrices into a truncated SVD and remainder as $\mathbf{Y}\boldsymbol{\Sigma}^{-1/2} = \mathbf{U}_{(k)}\mathbf{D}_{(k)}\mathbf{V}_{(k)}^\top + \mathbf{U}_{(-k)}\mathbf{D}_{(-k)}\mathbf{V}_{(-k)}^\top$, and observe that $\mathbf{U}^\top\mathbf{U} = \mathbf{I}$ implies $\mathbf{U}_{(k)}^\top\mathbf{U}_{(k)} = \mathbf{I}_{k\times k}$ and $\mathbf{U}_{(k)}^\top\mathbf{U}_{(-k)} = \mathbf{0}_{k\times n-k}$. In APEX, we use the expression $\hat{\mathbf{M}}_{(k)} = \mathbf{U}_{(k)}\mathbf{U}_{(k)}^\top\mathbf{Y}$ for computational convenience when $n < p$, without explicitly calculating the right singular vectors. The time complexity to calculate $\mathbf{U}_{(k)}$ is $O(n^2k)$, and to calculate $\hat{\mathbf{M}}_{(k)}$ is $O(pnk)$, giving an overall complexity of $O(n^2k + pnk)$. For $n > p$ (for example, in biobank-scale data sets with $n >$ 100,000), it is preferable calculate only the right singular vectors $\mathbf{V}_{(k)}$ and $\hat{\mathbf{M}}_{(k)} = \mathbf{Y}\boldsymbol{\Sigma}^{-1/2}\mathbf{V}_{(k)}\mathbf{V}_{(k)}^\top\boldsymbol{\Sigma}^{1/2}$, which together has time complexity $O(p^2k + pnk)$. Using this adaptive approach to calculate either the left or right singular vectors, the time complexity is $O\left\{\min(n,p)^2k + pnk\right\}$.

We additionally allow an inverse Wishart prior distribution for $\mathbf{\Sigma}$ with degree of freedom $\nu$ and scale matrix $\mathbf{I}\eta^2$, such that the updates $\tilde{\sigma}_j^2 = \frac{\eta^2 + \sum_{i=1}^n (Y_{ij} - M_{ij})^2}{\nu + p + 1 + n}$ maximize $\log \mathcal{L}_\pi(\mathbf{\Sigma}, \mathbf{M}) = \log \mathcal{L}(\mathbf{\Sigma}, \mathbf{M}) + \log \pi(\mathbf{\Sigma})$, where the prior likelihood (omitting constants) is

$$\log \pi(\mathbf{\Sigma}) = \frac{\nu}{2} \log |\mathbf{I}\eta^2| - \frac{\nu + p + 1}{2} \log |\mathbf{\Sigma}| - \frac{1}{2} \mathrm{tr}(\mathbf{I}\eta^2 \mathbf{\Sigma}^{-1}). \tag{4}$$

We parameterize $\nu$ and $\eta^2$ as $\mu_0 = \frac{\nu+p+1}{\nu+p+1+n}$ and $\sigma_0^2 = \frac{1}{\nu+p+1}\eta^2$, so that the updates become $\tilde{\sigma}_j^2 = \mu_0 \sigma_0^2 + (1 - \mu_0)\hat{\sigma}_j^2$. Here, the prior is improper for $\mu_0 \leq \frac{2p}{2p+n}$ (equivalently, $\nu \leq p - 1$).

Common factor variables inferred using the above procedure can be included as covariates in xQTL association analysis. This can reduce technical confounding and increase power by capturing residual variation, particularly in *cis* analysis. We note that inferred factors capturing genuine biological variation may also mask true *trans* signals if included as covariates in *trans* analysis. In *cis* or *trans* association analysis, factor covariates can be modeled either as fixed effects (using OLS, or REML when a genetic relatedness matrix is included in the model) or random effects (using REML; Section 0.3). For random effects, the resulting random-effect covariance matrix has rank $k < n$ (where $k$ is the number of inferred factors), which enables efficient computation for variance component estimation and LMM association tests.

#### 0.3 Efficient Variance Component Estimation for xQTL Studies

##### 0.3.1 The Mixed Model Likelihood Function

This section describes procedures in APEX for restricted maximum likelihood (REML) variance component estimation to account for familial relatedness in xQTL studies. We begin with the model in (1) for a single study and trait, $\boldsymbol{Y} = \mathbf{C}\boldsymbol{\zeta} + \mathbf{G}\boldsymbol{\beta} + \mathbf{Z}\boldsymbol{b} + \boldsymbol{\varepsilon}$, and define $\mathbf{\Omega} = \mathrm{Var}(\mathbf{Z}\boldsymbol{b} + \boldsymbol{\varepsilon}) = \mathbf{K}\tau^2 + \mathbf{I}\sigma^2$ as the total variance of $\boldsymbol{Y}$, where $\mathbf{K} = \mathbf{Z}\mathbf{Z}^\top$. For ease of exposition, we define $\mathbf{X} = (\mathbf{C}, \mathbf{G})$ and let $\boldsymbol{\alpha} = (\boldsymbol{\zeta}^\top, \boldsymbol{\beta}^\top)^\top$. In practice, APEX estimates variance components $\tau^2$ and $\sigma^2$ once for each gene under the null hypothesis $H_0 : \boldsymbol{\beta} = \mathbf{0}$ to calculate score statistics, similar to many LMM methods for genome-wide association studies (GWAS) (Lippert et al., 2011; Svishcheva et al., 2012; Loh et al., 2015; Zhou et al., 2018; Chen et al., 2019; Chen and Conomos, 2020). When evaluating the likelihood function to estimate variance components, the columns of $\mathbf{X}$ for which $\boldsymbol{\alpha}$ is fixed at 0 are simply excluded.

The REML log-likelihood function is

$$\ell_R(\tau^2, \sigma^2, \boldsymbol{\alpha}) = -\frac{1}{2} \log |\mathbf{\Omega}| - \frac{1}{2} \log |\mathbf{X}^\top \mathbf{\Omega}^{-1} \mathbf{X}| - \frac{1}{2} (\boldsymbol{Y} - \mathbf{X}\boldsymbol{\alpha})^\top \mathbf{\Omega}^{-1} (\boldsymbol{Y} - \mathbf{X}\boldsymbol{\alpha}) \tag{5}$$

Now we will re-write the likelihood to compute the MLEs more efficiently, and re-parameterize the variance components as $\sigma^2$ and $\theta := \tau^2/\sigma^2$, as has been described in several LMM methods in GWAS (Kang et al., 2010; Lippert et al., 2011; Zhou and Stephens, 2012). We decompose the matrix $\mathbf{K} = \mathbf{Q}\mathbf{\Lambda}\mathbf{Q}^\top$, where $\mathbf{\Lambda}$ is the diagonal matrix of eigenvalues and $\mathbf{Q}$ is the matrix of eigenvectors with $\mathbf{Q}^\top\mathbf{Q} = \mathbf{I}$. Define the diagonal matrix $\mathbf{D}_\theta = \mathbf{I} + \theta\mathbf{\Lambda}$, so that we can write $\mathbf{\Omega} = \sigma^2 \mathbf{Q}\mathbf{D}_\theta\mathbf{Q}^\top$. These procedures are further expedited for structured $\mathbf{K}$. When $\mathbf{K}$ is a block-diagonal GRM, $\mathbf{Q}$ is sparse and block-diagonal, with a diagonal block for unrelated individuals. When $\mathbf{K} = \mathbf{Z}\mathbf{Z}^\top$ for $\mathbf{Z}$ with $k$ columns (for example, $\mathbf{Z}$ are common factor covariates), we calculate only the $k$ eigenvectors $\mathbf{Q}_1$ and eigenvalues $\mathbf{\Lambda}_1$ with non-zero eigenvalues. Below, we separately detail LMM computations for (a) full-rank matrices $\mathbf{K}$, and (b) for low-rank matrices $\mathbf{K}$.

For full-rank $\mathbf{K}$, we premultiply $\mathbf{Q}^\top$ to the covariate and molecular trait matrices, and define these quantities as $\mathbf{X}_* = \mathbf{Q}^\top\mathbf{X}$ and $\boldsymbol{Y}_* = \mathbf{Q}^\top\boldsymbol{Y}$. We can write the re-parameterized REML log-likelihood as

$$\ell_R(\theta, \sigma^2, \boldsymbol{\alpha}) = -\frac{n-\nu}{2}\log\sigma^2 - \frac{1}{2}\log|\mathbf{D}_\theta| - \frac{1}{2}\log|\mathbf{X}_*^\top\mathbf{D}_\theta^{-1}\mathbf{X}_*| - \frac{1}{2\sigma^2}(\boldsymbol{Y}_* - \mathbf{X}_*\boldsymbol{\alpha})^\top\mathbf{D}_\theta^{-1}(\boldsymbol{Y}_* - \mathbf{X}_*\boldsymbol{\alpha}), \tag{6}$$

where $n$ is the sample size and $\nu$ is the rank of $\mathbf{X}$. When the genetic effects $\boldsymbol{\beta}$ are fixed to $\mathbf{0}$, we have $\nu = m$, where $m$ is the number of technical covariates including the intercept. Differentiating $\ell_R$ with respect to $\sigma^2$ and $\boldsymbol{\alpha}$ gives closed forms for the (restricted) maximum likelihood estimates (MLEs):

$$\hat{\boldsymbol{\alpha}}(\theta) = [\mathbf{X}_*^\top\mathbf{D}_\theta^{-1}\mathbf{X}_*]^{-1}\mathbf{X}_*^\top\mathbf{D}_\theta^{-1}\boldsymbol{Y}_*, \tag{7}$$

$$\hat{\sigma}^2(\theta, \boldsymbol{\alpha}) = \frac{1}{n-\nu}(\boldsymbol{Y}_* - \mathbf{X}_*\boldsymbol{\alpha})^\top\mathbf{D}_\theta^{-1}(\boldsymbol{Y}_* - \mathbf{X}_*\boldsymbol{\alpha}), \tag{8}$$

$$\hat{\sigma}^2_{\boldsymbol{\alpha}=\hat{\boldsymbol{\alpha}}}(\theta) = \frac{1}{n-\nu}\boldsymbol{Y}_L^\top\mathbf{A}_\theta\boldsymbol{Y}_L, \tag{9}$$

$$\mathbf{A}_\theta := \mathbf{D}_\theta^{-1} - \mathbf{D}_\theta^{-1}\mathbf{X}_*[\mathbf{X}_*^\top\mathbf{D}_\theta^{-1}\mathbf{X}_*]^{-1}\mathbf{X}_*^\top\mathbf{D}_\theta^{-1}. \tag{10}$$

Note that $\hat{\boldsymbol{\alpha}}(\theta)$ is the best linear unbiased estimate (BLUE) solution to the mixed model equations (Henderson, 1973).

Plugging in the above expressions, we obtain a closed-form profile likelihood that is only a function of $\theta$, which we can maximize to obtain its MLE $\hat{\theta}$:

$$l_R(\theta) := \ell_R(\sigma^2 = \hat{\sigma}^2, \theta, \boldsymbol{\alpha} = \hat{\boldsymbol{\alpha}}) \tag{11}$$

$$= -\frac{1}{2}\left\{(n-\nu)\log\hat{\sigma}^2 + \log|\mathbf{X}_*^\top\mathbf{D}_\theta^{-1}\mathbf{X}_*| + \log|\mathbf{D}_\theta| + \frac{1}{\hat{\sigma}^2}(\boldsymbol{Y}_* - \mathbf{X}_*\hat{\boldsymbol{\alpha}})^\top\mathbf{D}_\theta^{-1}(\boldsymbol{Y}_* - \mathbf{X}_*\hat{\boldsymbol{\alpha}})\right\} \tag{12}$$

$$= -\frac{1}{2}\left\{(n-\nu)\log\left(\boldsymbol{Y}_*^\top\mathbf{A}_\theta\boldsymbol{Y}_*\right) + \log|\mathbf{X}_*^\top\mathbf{D}_\theta^{-1}\mathbf{X}_*| + \log|\mathbf{D}_\theta|\right\} + \text{Constant}. \tag{13}$$

The above 1-dimensional profile likelihood has been extensively used to efficiently fit LMMs in GWAS (Kang et al., 2010; Zhou and Stephens, 2012; Jiang et al., 2019), and follows earlier work on variance component estimation (Patterson and Thompson, 1971; Harville, 1974).

For low-rank $\mathbf{K} = \mathbf{Z}\mathbf{Z}^\top$, we have $\boldsymbol{\Omega} = \sigma^2\left(\mathbf{I} + \theta\mathbf{Q}_1\mathbf{\Lambda}_1\mathbf{Q}_1^\top\right)$ and $\boldsymbol{\Omega}^{-1} = \sigma^{-2}\left(\mathbf{I} - \mathbf{Q}_1\mathbf{W}_\theta\mathbf{Q}_1^\top\right)$, where $\mathbf{W}_\theta = \theta\mathbf{\Lambda}_1\left(\mathbf{I} + \theta\mathbf{\Lambda}_1\right)^{-1}$. Then, we can similarly write the likelihood as

$$\ell_R(\theta,\sigma^2,\boldsymbol{\alpha}) = -\frac{n-\nu}{2}\log\sigma^2 - \frac{1}{2}\log\frac{\left|\mathbf{X}^\top\left(\mathbf{I}-\mathbf{Q}_1\mathbf{W}_\theta\mathbf{Q}_1^\top\right)\mathbf{X}\right|}{|\mathbf{I}-\mathbf{W}_\theta|} - \frac{1}{2\sigma^2}(\boldsymbol{Y}-\mathbf{X}\boldsymbol{\alpha})^\top\left(\mathbf{I}-\mathbf{Q}_1\mathbf{W}_\theta\mathbf{Q}_1^\top\right)(\boldsymbol{Y}-\mathbf{X}\boldsymbol{\alpha}). \quad (14)$$

In this case, we precompute the terms $\mathbf{X}_{*1} = \mathbf{Q}_1^\top\mathbf{X}$ and $\boldsymbol{Y}_{*1} = \mathbf{Q}_1^\top\boldsymbol{Y}$, $\mathbf{X}^\top\mathbf{X}$ and $\mathbf{X}^\top\boldsymbol{Y}$ to expedite computation. Then, the MLEs can be written

$$\hat{\boldsymbol{\alpha}}(\theta) = [\mathbf{X}^\top\mathbf{X} - \mathbf{X}_{*1}^\top\mathbf{W}_\theta\mathbf{X}_{*1}]^{-1}\left(\mathbf{X}^\top\boldsymbol{Y} - \mathbf{X}_{*1}^\top\mathbf{W}_\theta\boldsymbol{Y}_{*1}\right), \quad (15)$$

$$\hat{\sigma}^2_{\boldsymbol{\alpha}=\hat{\boldsymbol{\alpha}}}(\theta) = \frac{1}{n-\nu}\left\{\boldsymbol{Y}^\top\boldsymbol{Y} - \boldsymbol{Y}_{*1}^\top\mathbf{W}_\theta\boldsymbol{Y}_{*1} - \left(\boldsymbol{Y}^\top\mathbf{X} - \boldsymbol{Y}_{*1}^\top\mathbf{W}_\theta\mathbf{X}_{*1}\right)\hat{\boldsymbol{\alpha}}(\theta)\right\}, \quad (16)$$

which gives a similar 1-dimensional profile likelihood,

$$l_R(\theta) = -\frac{n-\nu}{2}\log\left\{\boldsymbol{Y}^\top\boldsymbol{Y} - \boldsymbol{Y}_{*1}^\top\mathbf{W}_\theta\boldsymbol{Y}_{*1} - \left(\boldsymbol{Y}^\top\mathbf{X} - \boldsymbol{Y}_{*1}^\top\mathbf{W}_\theta\mathbf{X}_{*1}\right)\hat{\boldsymbol{\alpha}}(\theta)\right\} + \frac{1}{2}\log|\mathbf{I}-\mathbf{W}_\theta| - \frac{1}{2}\log|\mathbf{X}^\top\mathbf{X} - \mathbf{X}_{*1}^\top\mathbf{W}_\theta\mathbf{X}_{*1}| + \text{Constant}. \quad (17)$$

##### 0.3.2 Efficient LMM Computation with Structured GRMs

When eFA or ePC covariates are modeled as random effects in LMMs, the variance-covariance matrix of random effects $\mathbf{K} = \mathbf{Z}\mathbf{Z}^\top$ has rank $k < n$, where $k$ is the number of eFA or ePC covariates (columns of $\mathbf{Z}$). APEX exploits the low-rank structure of these matrices to expedite computation; similar strategies have been used low-rank GRMs constructed from a sparse set genetic variants (Lippert et al., 2011). APEX also supports sparse and block-diagonal GRMs to account for familial relatedness (Jiang et al., 2019; Gogarten et al., 2019).

###### Time Complexity of Variance Component Estimation

Here, we show time complexity of variance component estimation for (a) full-rank covariance matrix $\mathbf{K}$ and (b) for low-rank $\mathbf{K}$. For full-rank $\mathbf{K}$, we first must decompose $\mathbf{K} = \mathbf{Q}\boldsymbol{\Lambda}\mathbf{Q}^\top$, which is in general $O(n^3)$. However, when $\mathbf{K}$ is sparse with a block diagonal structure the decomposition at most $O(n\overline{b^3})$ where $\overline{b^3}$ is the average cubed familial block size. For instance, in a sample consisting of unrelated trios, $\overline{b^3} = 9$. In a sample where $pn$ samples are related in blocks of size $b$ and the remaining $(1-p)n$ are unrelated, $\overline{b^3} = pb^3 + (1-p)$, and the decomposition is at most $O(npb^3)$. Next, we estimate the variance components for each of the molecular traits. To evaluate the profile likelihood $l_R(\theta)$, we must calculate:

1. $\boldsymbol{Y}_*^\top\mathbf{D}_\theta^{-1}\boldsymbol{Y}_*$, which is $O(n)$ as $\mathbf{D}_\theta^{-1}$ is diagonal.

2. $\log|\mathbf{X}_*^\top\mathbf{D}_\theta^{-1}\mathbf{X}_*|$, which is $O(m^2n)$ where $m$ is the number of covariates, since $m < n$.

3. $[\mathbf{X}_*^\top\mathbf{D}_\theta^{-1}\mathbf{X}_*]^{-1}\mathbf{X}_*^\top\mathbf{D}_\theta^{-1}\boldsymbol{Y}_*$, which is $O(m^2n)$, since $m < n$.

Therefore, the time complexity of variance component estimation with full-rank $\mathbf{K}$ is $O(m^2n)$ for a single molecular trait and $O(pm^2n)$ across all traits where $p$ is the number of molecular traits, $m$ the number of covariates, and $n$ the sample size.

For a low-rank matrix $\mathbf{K} = \mathbf{Z}\mathbf{Z}^\top$ where $\mathbf{Z}$ is full rank with $k$ columns, we first pre-calculate the terms $\mathbf{X}_{*1} = \mathbf{Q}_1^\top \mathbf{X}$ and $\boldsymbol{Y}_{*1} = \mathbf{Q}_1^\top \boldsymbol{Y}$, $\mathbf{X}^\top\mathbf{X}$ and $\mathbf{X}^\top \boldsymbol{Y}$. Then, to evaluate the profile likelihood $l_R(\theta)$ for a given trait, we must calculate:

1. $\mathbf{X}_{*1}^\top \mathbf{W}_\theta \boldsymbol{Y}_{*1}$, which is $O(mk)$.
2. $\log |\mathbf{X}^\top\mathbf{X} - \mathbf{X}_{*1}^\top \mathbf{W}_\theta \mathbf{X}_{*1}|$, which is $O(m^3 + m^2 k)$.
3. $[\mathbf{X}^\top\mathbf{X} - \mathbf{X}_{*1}^\top \mathbf{W}_\theta \mathbf{X}_{*1}]^{-1} \left(\mathbf{X}^\top \boldsymbol{Y} - \mathbf{X}_{*1}^\top \mathbf{W}_\theta \boldsymbol{Y}_{*1}\right)$, which is $O(m^3)$.

Therefore, the time complexity of variance component estimation with a rank-$k$ matrix $\mathbf{K}$ is $O(m^3 + m^2 k)$ for a single molecular trait, and $O(pm^3 + pm^2 k)$ across all traits where $p$ is the number of molecular traits, $m$ the number of covariates, and $k$ the rank of $\mathbf{K}$.

In both cases, we use Brent's derivative-free algorithm (Brent, 1973) to optimize the 1-dimensional likelihood as described in (Kang et al., 2010); in practice, we find that the algorithm converges quickly. In general, we have found that the time required to estimate variance components is trivial relative to singe-variant analysis.

#### 0.4 xQTL Association Tests in the Linear Mixed Model

Here we describe association test statistics under the linear mixed model. The score statistics for a given trait are

$$\boldsymbol{U}_\beta := \left.\frac{\partial \ell_R}{\partial \boldsymbol{\beta}}\right|_{\boldsymbol{\zeta}=\hat{\boldsymbol{\zeta}}, \tau^2=\hat{\tau}^2, \sigma^2=\hat{\sigma}^2, \boldsymbol{\beta}=\mathbf{0}} = \mathbf{G}^\top \mathbf{P} \boldsymbol{Y}, \tag{18}$$

and

$$\mathbf{V}_\beta = \hat{\text{Var}}\left(\boldsymbol{U} \middle| \hat{\boldsymbol{\zeta}}, \hat{\tau}^2, \hat{\sigma}^2\right) = \mathbf{G}^\top \mathbf{P} \mathbf{G}, \tag{19}$$

where $\mathbf{P} := \hat{\boldsymbol{\Omega}}^{-1}[\mathbf{I} - \mathbf{C}(\mathbf{C}^\top \hat{\boldsymbol{\Omega}}^{-1} \mathbf{C})^{-1} \mathbf{C}^\top \hat{\boldsymbol{\Omega}}^{-1}]$. The matrix $\mathbf{V}_\beta$ is the Fisher information for $\boldsymbol{\beta}$ conditional on the covariate effect estimates $\hat{\boldsymbol{\zeta}} = [\mathbf{C}^\top \hat{\boldsymbol{\Omega}}^{-1} \mathbf{C}]^{-1} \mathbf{C}^\top \hat{\boldsymbol{\Omega}}^{-1} \boldsymbol{Y}$ and variance component MLEs $\hat{\sigma}^2$ and $\hat{\tau}^2$.

##### 0.4.1 Approximate $F$ Tests for Genetic Association in the LMM

Here we describe approximately $F$ and $t$ distributed LMM association test statistics implemented in APEX. These tests were selected with two features in mind. First, they provide accurate p-values with small sample sizes and/or large numbers of technical covariates (e.g., PEER factors) by explicitly accounting for the residual degrees of freedom. Second, they can be calculated efficiently with large numbers of markers and traits, and can be written simply as functions of the single-variant score statistics $\boldsymbol{U}_\beta$ and variance matrix $\mathbf{V}_\beta$ as defined above.

We propose a modification of a widely-used approximate $F$ test statistic for fixed effect hypotheses of the form $H_0 : \mathbf{L}\boldsymbol{\beta} = \mathbf{0}$ in LMMs (Kuznetsova, Brockhoff, and Christensen, 2017; Halekoh and Højsgaard, 2014; Giesbrecht and Burns, 1985; Hrong-Tai Fai and Cornelius, 1996). Recall that the MLEs $\hat{\boldsymbol{\beta}}(\theta)$ and $\hat{\sigma}^2(\theta)$ have closed form given a fixed value of the nuisance parameter $\theta = \tau^2/\sigma^2$. Let $\hat{\theta}_0$ denote the MLE of $\theta$ under the null hypothesis $\mathbf{L}\boldsymbol{\beta} = \mathbf{0}$. The APEX approximate $F$ test is

$$\tilde{F} = \frac{\left(\mathbf{L}\hat{\boldsymbol{\beta}}(\hat{\theta}_0)\right)^\top \left(\mathbf{L}\hat{\text{Var}}[\hat{\boldsymbol{\beta}}(\hat{\theta}_0)]\mathbf{L}^\top\right)^{-1} \left(\mathbf{L}\hat{\boldsymbol{\beta}}(\hat{\theta}_0)\right)}{\text{rank}(\mathbf{L})}, \tag{20}$$

which is asymptotically equivalent under $H_0$ to the standard LMM $F$ test using $\hat{\theta}$ rather than $\hat{\theta}_0$.While $\tilde{F}$ is clearly a Wald-type test, it also has the appealing property that it is numerically equivalent to the OLS $F$ test when $\hat{\theta}_0 = 0$.

For single-variant association tests, the corresponding t-test statistic is given by

$$\tilde{t} = \frac{\hat{\beta}_{\hat{\theta}_0}}{SE(\hat{\beta}_{\hat{\theta}_0})}, \tag{21}$$

where the slope $\hat{\beta}_{\hat{\theta}_0} = \frac{\boldsymbol{G}^\top \mathbf{A}_{\hat{\theta}_0} \boldsymbol{Y}}{\boldsymbol{G}^\top \mathbf{A}_{\hat{\theta}_0} \boldsymbol{G}}$, standard error $SE(\hat{\beta}_{\hat{\theta}_0}) = \sqrt{\hat{\sigma}^2_{\hat{\theta}_0} \left(\boldsymbol{G}^\top \mathbf{A}_{\hat{\theta}_0} \boldsymbol{G}\right)^{-1}}$, and the matrix

$$\mathbf{A}_{\hat{\theta}_0} = (\mathbf{K}\hat{\theta}_0 + \mathbf{I})^{-1} \left(\mathbf{I} - \mathbf{C}[\mathbf{C}^\top (\mathbf{K}\hat{\theta}_0 + \mathbf{I})^{-1}\mathbf{C}]^{-1}\mathbf{C}^\top (\mathbf{K}\hat{\theta}_0 + \mathbf{I})^{-1}\right). \tag{22}$$

While $\theta := \tau^2/\sigma^2$ is estimated under the null hypothesis $\beta = 0$, note that we update the residual variance component estimate $\sigma^2$ to calculate the standard error ( holding constant the ratio $\tau^2/\sigma^2 = \theta$). Specifically, $\hat{\sigma}^2_{\hat{\theta}_0} = \frac{1}{n-m-1}\boldsymbol{Y}^\top \mathbf{A}^*_{\hat{\theta}_0} \boldsymbol{Y}$, where $\mathbf{A}^*_{\hat{\theta}_0}$ identical to $\mathbf{A}_{\hat{\theta}_0}$ with $\mathbf{C}$ replaced by $[\mathbf{C}, \boldsymbol{G}]$, which is the closed-form MLE for $\sigma^2$ as described above.

**Relationship between the score and approximate $t$ test statistics.** Here we describe the relationship between the approximate $t$ test and the score test, which is widely used in GWAS mixed models as it only requires MLEs evaluated under the null hypothesis, which can be calculated once and re-used for single-variant association tests. The LMM score test is here given by

$$S = \frac{\boldsymbol{G}^\top \mathbf{A}_{\hat{\theta}_0} \boldsymbol{Y}}{\sqrt{\hat{\sigma}^2_{H_0} \boldsymbol{G}^\top \mathbf{A}_{\hat{\theta}_0} \boldsymbol{G}}}, \tag{23}$$

where $\hat{\sigma}^2_{H_0} = \boldsymbol{Y}^\top \mathbf{A}_{\hat{\theta}_0} \boldsymbol{Y}/(n-m)$. Under the null hypothesis $\beta = 0$, $S$ asymptotically follows the standard normal distribution. The approximate $t$ test statistic $\tilde{t}$ can be related to the score test $S$ by the equation $\hat{\sigma}_{\hat{\theta}_0}\tilde{t} = \hat{\sigma}_{H_0} S$, where recall that $\hat{\sigma}^2_{\hat{\theta}_0} = \frac{1}{n-m-1}\boldsymbol{Y}^\top \mathbf{A}^*_{\hat{\theta}_0} \boldsymbol{Y}$. By expanding $\mathbf{A}^*_{\hat{\theta}_0}$, it can be shown that $\hat{\sigma}^2_{\hat{\theta}_0} = \frac{n-m}{n-m-1}\hat{\sigma}^2_{H_0} + \frac{1}{n-m-1}\hat{\sigma}^2_{H_0} S^2$, which implies that $\tilde{t}^2 > S^2$ whenever $S^2 > 1$. Importantly, however, the significance of $S^2$ is often evaluated using the asymptotic normal distribution in finite samples. To calculate p-values for $\tilde{t}^2$, we explicitly account for the residual degrees of freedom $\nu$, which is roughly $n - m - 1$ where $n$ is the sample size and $m$ is the number of covariates, and can be approximated more accurately by the Satterthwaite or Kenward-Roger methods (Welch, 1947; Satterthwaite, 1946; Kenward and Roger, 1997; Kuznetsova, Brockhoff, and Christensen, 2017). The Satterthwaite approximation in this case is $\tilde{\nu} = \frac{2SE(\hat{\beta}_{\hat{\theta}_0})^2}{\hat{\text{Var}}[SE(\hat{\beta}_{\hat{\theta}_0})^2]}$,where the denominator can be approximated using the delta method.

##### 0.4.2 Efficient Computation of LMM Single-Variant Score Test Denominators

Here we describe procedures to efficiently calculate the denominator terms $V_{jt} = \boldsymbol{G}_j^\top \mathbf{P}_t \boldsymbol{G}_j$ across traits $t = 1, 2, ..., p$ and variants $j = 1, 2, ..., q$. Because $\mathbf{P}_t \boldsymbol{Y}_t$ can be calculated once for each trait $t$ and recycled, the time complexity to calculate numerator terms $U_{jt}$ is $O(n)$ by dense product or $O(K_j)$ by sparse product where $K_j$ is the number of non-zero entries in $\boldsymbol{G}_j$. However, the denominator terms $V_{jt}$ are more difficult, as direct calculation would be $O(n^2)$ for each variant-trait pair by dense product or $O(nK_j)$ by sparse product. Moreover, this is particularly challenging for QTL analysis, as $V_{jt}$ must be calculated across tens of thousands of molecular traits for each variant.

Many methods have addressed this problem using the GRAMMAR-GAMMA approximation, essentially $V_{jt} = \boldsymbol{G}_j^\top \mathbf{P}_t \boldsymbol{G}_j \approx \gamma \boldsymbol{G}_j^\top (\mathbf{I} - \tilde{\mathbf{H}})\boldsymbol{G}_j$ where $\tilde{\mathbf{H}}$ is the ordinary least squares (OLS) hat matrix calculated from covariates $\mathbf{C}$ or a subset thereof (e.g., only the intercept column) (Svishcheva et al., 2012; Loh et al., 2015; Zhou et al., 2018; Jiang et al., 2019). This approximation is expected to perform well when genotypes are uncorrelated with fixed-effect covariates $\mathbf{C}$ and/or random-effect covariates $\mathbf{Z}$. In APEX, we take a different approach to accurately approximate each $V_{jt}$ under arbitrary correlations between $\mathbf{G}$, $\mathbf{C}$, and $\mathbf{Z}$ for common and rare variants and in both small and large sample sizes. First, we notice $\mathbf{P}_t$ can be written as $\mathbf{P}_t = \hat{\sigma}_t^2 \mathbf{A}(\hat{\theta}_t)$ where $\mathbf{A}(\theta)$ has a common form across traits and depends only on the ratio $\theta = \tau^2/\sigma^2$. Next, we define the function $f_j(\theta) = \boldsymbol{G}_j^\top \mathbf{A}(\theta)\boldsymbol{G}_j$, so that each $V_{jt}$ can be written $V_{jt} = \hat{\sigma}_t^2 f_j(\theta)$. We evaluate the function $f_j()$ by expanding

$$f_j(\theta) = \boldsymbol{G}_j^\top \boldsymbol{G}_j - \boldsymbol{G}_j^\top \tilde{\mathbf{Q}}\boldsymbol{\Psi}(\theta)\tilde{\mathbf{Q}}^\top \boldsymbol{G}_j - \boldsymbol{r}_j(\theta)^\top \left(\mathbf{C}^\top\mathbf{C} - \mathbf{C}^\top\tilde{\mathbf{Q}}\boldsymbol{\Psi}(\theta)\tilde{\mathbf{Q}}^\top\mathbf{C}\right)^{-1} \boldsymbol{r}_j(\theta) \tag{24}$$

where $\boldsymbol{r}_j(\theta) = \mathbf{C}^\top \boldsymbol{G}_j - \mathbf{C}^\top\tilde{\mathbf{Q}}\boldsymbol{\Psi}(\theta)\tilde{\mathbf{Q}}^\top \boldsymbol{G}_j$. When $\mathbf{K}$ is full rank, $\tilde{\mathbf{Q}}$ are the eigenvectors with eigenvalues $\neq 1$ and $\boldsymbol{\Psi}(\theta) = \text{diag}\left(\frac{(\lambda_k - 1)\theta}{1+(\lambda_k-1)\theta}\right)$. When $\mathbf{K}$ is low-rank, $\tilde{\mathbf{Q}}$ are the eigenvectors with eigenvalue $\neq 0$ and $\boldsymbol{\Psi}(\theta) = \text{diag}\left(\frac{\lambda_k\theta}{1+\lambda_k\theta}\right)$. These computations are expedited by pre-computing and storing the re-used terms in $f_j$.

Each function $f_j(\theta)$ is smooth and finite for all $\theta \in [0, \infty]$, and we approximate $f_j(\theta)$ using a spline interpolation with a fixed grid of sample points $\theta_s \in \{0, \theta_2, ..., \theta_{m-1}, \infty\}$. In practice, we find that simply taking $\theta_s \in \{0, 1, \infty\}$ provides highly accurate approximations. Overall, calculating $f_j$ for all variant $j = 1, 2, ...., q$ is $O(nms + nmq)$, where $s = \overline{b^2}$ for block-diagonal $\mathbf{K}$ (the mean squared block size) or $s = k$ for rank-$k$ $\mathbf{K}$, $m$ is the number of covariates, and $n$ the sample size. Once completed, each $V_{jt}$ can be calculated in $O(1)$. This reduces the overall time complexity of association testing from $O(nms+nmpq)$ to $O(nms+nmq+npq)$, where $n$ is the sample size, $m$ is the number of technical covariates, $p$ is the number of traits, and $q$ is the number of genetic variants. For a single trait, this procedure is superfluous; however, it is economical for 4 or more traits, and vastly reduces computation time for *trans*-xQTL analysis.

#### 0.5 Secondary xQTL Discovery Algorithms

Single-variant association analysis aims to detect and estimate associations between individual genetic variants and traits, which tend to occur in clusters along a chromosome due to linkage disequilibrium (LD). Also of interest is whether single-variant association signals are driven by one or multiple causal variants, and the joint effects of multiple correlated variants. This section describes methods to detect multiple correlated genetic associations in APEX. Here, we use a Frequentist multiple regression framework; however, we note that APEX xQTL summary data files support a wider range of multiple-variant association methods, including elastic net and Bayesian sparse regression.

We again use the model $\boldsymbol{Y} = \mathbf{C}\boldsymbol{\zeta} + \mathbf{G}\boldsymbol{\beta} + \mathbf{Z}\boldsymbol{b} + \boldsymbol{\varepsilon}$, where $\boldsymbol{b} \sim \mathcal{N}(\mathbf{0}, \mathbf{I}\tau^2)$ and $\boldsymbol{\varepsilon} \sim \mathcal{N}(\mathbf{0}, \mathbf{I}\sigma^2)$. Similar to single-variant analysis, we update the MLE of $\sigma^2$ in each candidate regression model while holding the ratio $\theta = \tau^2/\sigma^2$ constant at its MLE $\hat{\theta}_0$ under the global null hypothesis $H_0 : \boldsymbol{\beta} = \mathbf{0}$. In APEX, we use stepwise regression procedures to select a sequence of associated variants $s_1, s_2, ..., s_M$, and use a stopping rule to choose the final number of tested signals $M$ while maintaining family-wise error rate FWER $< \alpha$ for some specified $\alpha$, accounting for the total number of variants tested and LD between them.

The basic stepwise algorithm proceeds as follows for each iteration $i$ beginning at $i = 0$:

1. Calculate conditional association test statistics $\tilde{t}_{c,i}$ for each $c \in C_i$, conditioning on both technical covariates $\mathbf{C}$ and selected variants $\mathbf{G}_{S_s}$.

2. Calculate an omnibus p-value $p_s^O$ for the null hypothesis $H_0 : \beta_k = 0$ for all $k \in C_i$.

3. If $p_i^O < \alpha$, then add $c_i^* = \arg\max_{c \in C_i} \tilde{t}_{c,i}^2$ to the selected set $S_{i+1} \leftarrow S_i \cup \{c_i^*\}$, update the candidate set $C_{i+1} \leftarrow C_i \backslash \{c_i^*\}$, set $i \leftarrow i + 1$ and return to step 1. Otherwise, stop and set the final number of variants $M \leftarrow i$.

The global FWER is here given by $P(M > 0|\boldsymbol{\beta} = \mathbf{0})$. The above procedure clearly maintains global FWER $= P(p_0^O < \alpha|\boldsymbol{\beta} = \mathbf{0}) \leq \alpha$ if $p_0^O$ is a valid p-value. Also, $P(p_i^O < \alpha|\boldsymbol{\beta}_{C_i} = \mathbf{0}) \leq \alpha$ at each step $i$, where $\boldsymbol{\beta}_{C_i}$ denotes the effects of variants in the candidate set, if $p_i^O$ is a valid p-value. We calculate $p_i^O$ using the Cauchy p-value combination method (ACAT) (Liu and Xie, 2020), which provides valid p-values for arbitrary dependence structures, can be calculated efficiently across thousands of variants, and optionally incorporates functional prior weights.

APEX implements 4 extensions of the simple forward stepwise procedure described above. First, we calculate the variance inflation factor (VIF) at each iteration $i$ for all variants in the candidate set $C_i$ given technical covariates $\mathbf{C}$ and selected variant genotypes $\mathbf{G}_{S_i}$, and prune any variants with VIF larger than a specified threshold (4 by default) from the candidate set. Second, we add a backward step to remove selected variants that are no longer significant in the joint model. At each iteration $i$, we calculate the $t$ test p-values of each selected variant $s \in S_i$ in the model including $\mathbf{C}$ and $\mathbf{G}_{S_i}$, and drop least significant variant from $S_i$ if its p-value is greater than a specified threshold ($\alpha$ by default, or 1 to omit the backward step). Finally, we implemented these procedures in a form that requires only association summary statistics and variance-covariance data to allow multiple-variant meta-analysis in the absence of individual-level data. We additionally allow for heterogeneity in genetic effects across studies in meta-analysis by using modified conditional test statistics (estimating genetic effects separately within each study), which are described in the subsequent section.

#### 0.6 Single- and Multiple-Variant xQTL Meta-Analysis

##### Individual-Level Data

The table below defines the basic individual-level data matrices xQTL analysis and meta-analysis. Per-study matrices (column 1) are used to perform standard *cis* and *trans* xQTL analysis in a single study, and to generate summary statistics (defined in the subsequent section) for meta-analysis when individual-level data cannot be shared. Meta-analysis matrices (column 2), which combine individual-level matrices across studies, are defined for exposition only, and are not explicitly computed or stored.

Here, we assume that that same sets of molecular traits $\mathbf{Y}_s$ and genetic variants $\mathbf{G}_s$ are measured across all studies; in practice, APEX meta-analyzes the union of molecular traits across studies in which they were measured, and intersection of variants across studies. The set of technical covariates $\mathbf{C}_s$ may differ across studies; for example, genotype PCs or PEER factors are typically calculated separately within each study, and age and sex may be included as explicit covariates or implicitly captured by PEER factors (Price et al., 2006; Stegle et al., 2012; Consortium, 2017; Aguet et al., 2019). This is not problematic if the study-specific covariates $\mathbf{C}_s$ are each designed to capture a common set of unobserved confounders (for example, population structure and batch effects). Specifically, suppose that the covariates in each study capture a common set of unobserved confounders $\mathbf{C}_s^* \in \text{span}(\mathbf{C}_s)$, which implies that $\mathbf{C}_s^* = \mathbf{C}_s(\mathbf{C}_s^\top \mathbf{C}_s)^{-1}\mathbf{C}_s^\top \mathbf{C}_s^*$, and so $\mathbf{C}_s^* \boldsymbol{\psi} = \mathbf{C}_s \boldsymbol{\zeta}$ where $\boldsymbol{\zeta}_s = (\mathbf{C}_s^\top \mathbf{C}_s)^{-1}\mathbf{C}_s^\top \mathbf{C}_s^* \boldsymbol{\psi}$.

| | Per Study $i$ | Meta-Analysis |
|---|---|---|
| Trait Matrix | $\mathbf{Y}_s : n_s \times p_s$ | $\mathbf{Y}_\bullet = [\mathbf{Y}_1^\top, \mathbf{Y}_2^\top, ..., \mathbf{Y}_N^\top]^\top$ |
| Trait $t$ Covariance Matrix | $\boldsymbol{\Omega}_{st} : n_s \times n_s$ | $\boldsymbol{\Omega}_{\oplus t} = \bigoplus_{s=1}^N \boldsymbol{\Omega}_{st}$ |
| Technical Covariate Matrix | $\mathbf{C}_s : n_s \times m_s$ | $\mathbf{C}_\oplus = \bigoplus_{s=1}^N \mathbf{C}_s$ |
| Covariate Hat Matrix | $\mathbf{H}_s = \mathbf{C}_s(\mathbf{C}_s^\top \boldsymbol{\Omega}_s^{-1} \mathbf{C}_s)^{-1} \mathbf{C}_s^\top \boldsymbol{\Omega}_s^{-1}$ | $\mathbf{H}_\oplus = \bigoplus_{s=1}^N \mathbf{H}_s$ |
| Genotype Matrix, Homogeneous | $\mathbf{G}_s : n_s \times q_s$ | $\mathbf{G}_\bullet = [\mathbf{G}_1^\top, \mathbf{G}_2^\top, ..., \mathbf{G}_N^\top]^\top$ |
| Genotype Matrix, Heterogeneous | — | $\mathbf{G}_\oplus = \bigoplus_{s=1}^N \mathbf{G}_s$ |

Here and elsewhere, $\bigoplus$ denotes the direct sum, so that $\bigoplus_j \mathbf{X}_j = \text{diag}(\mathbf{X}_1, \mathbf{X}_2, ...)$ is a block-diagonal matrix with dimension given by the sums of the numbers of rows and columns of the input matrices. $\mathbf{I}$ denotes the identity matrix, $\mathbf{0}$ denotes the matrix of 0s, and $\mathbf{1}$ the column vector of 1s of implicit dimensions. We refer to the $k^{th}$ column of $\mathbf{G}_s$ as $\boldsymbol{G}_{s;k}$, and define $\mathbf{G}_{s;\boldsymbol{k}} = [\boldsymbol{G}_{s;k_1}, \boldsymbol{G}_{s;k_2}, ...]$ for subset of columns $\boldsymbol{k} = \{k_1, k_2, ...\}$.

##### Single-Variant Association Summary Statistics

Below we define summary association statistics as functions of individual-level data. These data are assumed to be shared across studies for meta-analysis, and are similar to summary statistics used in rare-variant meta-analysis toolkits (Feng et al., 2014; Zhan et al., 2016).

| | Per Study $i$ | Meta-Analysis |
|---|---|---|
| Score vector | $\boldsymbol{U}_s = \mathbf{G}_s^\top \boldsymbol{\Omega}_s^{-1}(\mathbf{I} - \mathbf{H}_s)\boldsymbol{Y}_s$ | $\boldsymbol{U}_\bullet = \sum_{s=1}^N \boldsymbol{U}_s$ |
| Variance-covariance matrix | $\mathbf{V}_s = \mathbf{G}_s^\top \boldsymbol{\Omega}_s^{-1}(\mathbf{I} - \mathbf{H}_s)\mathbf{G}_s$ | $\mathbf{V}_\bullet = \sum_{s=1}^N \mathbf{V}_s$ |
| Residual Sum of Squares | $S_s = \boldsymbol{Y}_s^\top \boldsymbol{\Omega}_s^{-1}(\mathbf{I} - \mathbf{H}_s)\boldsymbol{Y}_s$ | $S_\bullet = \sum_s S_s = \boldsymbol{Y}_\bullet^\top \boldsymbol{\Omega}_\oplus^{-1}(\mathbf{I} - \mathbf{H}_\oplus)\boldsymbol{Y}_\bullet$ |

##### Multiple Regression Meta-Analysis: Slopes and Score Statistics

The table below gives expression for the score statistic $U_j^*$, variance $V_j^*$, and residual sum of squares $S^*$ under the null hypothesis for a single variant $j$ in single-variant (marginal) meta-analysis, and multiple-variant meta-analysis with homogeneous or heterogeneous effects across studies. In the homogeneous-effect model, we assume the effect of each variant is constant across studies, whereas in the heterogeneous-effects model, we condition on variants separately within each study to account for differences in variant effects across studies. Expressions for each term in each model are given in the table below as functions of marginal single-variant summary statistics, although they can also be expressed in terms of the underlying individual-level data matrices.

**Conditional score, variance, and sum of squares from summary statistics.**

| | Marginal | Joint, homogeneous | Joint, heterogeneous (for a single study $s$) |
|---|---|---|---|
| Score statistic, $U_j^*$ | $U_{\bullet;j}$ | $U_{\bullet;j} - \boldsymbol{V}_{\bullet;\boldsymbol{k},j}^\top \mathbf{V}_{\bullet;\boldsymbol{k},\boldsymbol{k}}^{-1} \boldsymbol{U}_{\bullet;\boldsymbol{k}}$ | $U_{s;j} - \boldsymbol{V}_{s;\boldsymbol{k},j}^\top \mathbf{V}_{s;\boldsymbol{k},\boldsymbol{k}}^{-1} \boldsymbol{U}_{s;\boldsymbol{k}}$ |
| Variance of score, $V_j^*$ | $V_{\bullet;j,j}$ | $V_{\bullet;j,j} - \boldsymbol{V}_{\bullet;\boldsymbol{k},j}^\top \mathbf{V}_{\bullet;\boldsymbol{k},\boldsymbol{k}}^{-1} \boldsymbol{V}_{\bullet;\boldsymbol{k},j}$ | $V_{s;j,j} - \boldsymbol{V}_{s;\boldsymbol{k},j}^\top \mathbf{V}_{s;\boldsymbol{k},\boldsymbol{k}}^{-1} \boldsymbol{V}_{s;\boldsymbol{k},j}$ |
| Residual SS, $S^*$ | $S_\bullet$ | $S_\bullet - \boldsymbol{U}_{\bullet;\boldsymbol{k}}^\top \mathbf{V}_{\bullet;\boldsymbol{k},\boldsymbol{k}}^{-1} \boldsymbol{U}_{\bullet;\boldsymbol{k}}$ | $S_s - \boldsymbol{U}_{s;\boldsymbol{k}}^\top \mathbf{V}_{s;\boldsymbol{k},\boldsymbol{k}}^{-1} \boldsymbol{U}_{s;\boldsymbol{k}}$ |

Note that $S^*$ is defined as the sum of squared trait results adjusting only for technical covariates $\mathbf{C}$ and the set of conditioned variants $\boldsymbol{k}$, and not including the tested variant $j$. The additional sum of squares due to variant $j$ given $\mathbf{C}$ and any conditioned variants $\boldsymbol{k}$ is $(U_j^*)^2/V_j^*$, and the residual sum of squares including $j$ is $S^* - (U_j^*)^2/V_j^*$.

In each of the above models (3 columns), the regression slope for variant $j$ can be written as $\hat{\beta}_j^* = U_j^*/V_j^*$. For heterogeneous effects across studies (third column), we either separately estimate $\hat{\beta}_{s,j}^*$ in each study and test $H_0 : \hat{\beta}_{s,j}^* = 0 \forall s = 1, 2, ..N$ using an $F$ test, or estimate a combined slope across studies using $U_j^* = \sum_s U_{s,j}^*$ and $V_j^* = \sum_s V_{s,j}^*$ (allowing the effects of conditioned variants $\boldsymbol{k}$ to differ across studies); the test form can be specified in the APEX command line interface. These formula and procedures to calculate genotype regression coefficients are encapsulated by the following three rules:

**1. The slope and standard error for $\boldsymbol{G}$ in the model $Y = \mathbf{C}\boldsymbol{\zeta} + \boldsymbol{G}\beta + \boldsymbol{\epsilon}$, where $\boldsymbol{\epsilon} \sim \mathcal{N}(\mathbf{0}, \mathbf{I}\sigma^2)$, are:**

$$\hat{\beta} = \frac{U}{V} \qquad \text{and} \qquad se_{\hat{\beta}}^2 = \frac{1}{N-1}\left(\frac{S}{V} - \frac{U^2}{V^2}\right), \tag{25}$$

where $U = \sigma^{-2}\boldsymbol{G}^\top(\mathbf{I} - \mathbf{H})\boldsymbol{Y}$ is the score statistic, $V = \sigma^{-2}\boldsymbol{G}^\top(\mathbf{I} - \mathbf{H})\boldsymbol{G}$ the variance, $S = \sigma^{-2}\boldsymbol{Y}^\top(\mathbf{I} - \mathbf{H})\boldsymbol{Y}$ the sum of squared residuals (excluding $\boldsymbol{G}$), $\mathbf{H}$ is the hat matrix formed by $\mathbf{C}$, and $N = \text{length}(\boldsymbol{Y}) - \text{rank}(\mathbf{C})$. The $t$-test statistic $t_{\hat{\beta}} = \hat{\beta}/se_{\hat{\beta}}$ has $N - 1$ degrees of freedom. Also note that $\hat{\beta}$ and $se_{\hat{\beta}}^2$ do not depend on the parameter $\sigma^2$, which cancels out in both $U/V$ and $S/V$. Therefore, we can calculate the slopes by substituting $\sigma^2 = 1$ in the expressions for $U, V$ and $S$.

**2. To condition on additional covariates $\mathbf{G}_0$, update the slope and standard error for $\boldsymbol{G}$ as follows:**

$$U^{\text{new}} = U - \boldsymbol{V}_{G_0,G}^\top \mathbf{V}_{G_0}^{-1} \boldsymbol{U}_{G_0} \tag{26}$$

$$V^{\text{new}} = V - \boldsymbol{V}_{G_0,G}^\top \mathbf{V}_{G_0}^{-1} \boldsymbol{V}_{G_0,G} \tag{27}$$

$$S^{\text{new}} = S - \boldsymbol{U}_{G_0}^\top \mathbf{V}_{G_0}^{-1} \boldsymbol{U}_{G_0} \tag{28}$$

where $\boldsymbol{V}_{G_0,G} = \sigma^{-2}\mathbf{G}_0^\top(\mathbf{I} - \mathbf{H})\boldsymbol{G}$, $\mathbf{V}_{G_0} = \sigma^{-2}\mathbf{G}_0^\top(\mathbf{I} - \mathbf{H})\mathbf{G}_0$, and $\boldsymbol{U}_{G_0} = \sigma^{-2}\mathbf{G}_0^\top(\mathbf{I} - \mathbf{H})\boldsymbol{Y}$. The new slope and standard error for $\boldsymbol{G}$ conditioning on $\mathbf{G}_0$ in addition to $\mathbf{C}$ can be calculated again plugging $U^{\text{new}}$, $V^{\text{new}}$, $S^{\text{new}}$, and $N^{\text{new}} := N^{\text{new}} = N - \text{length}(\boldsymbol{U}_{G_0})$ into the expressions in 1.

Under the "heterogeneous-effect" model in meta-analysis, both $\mathbf{G}_0$ and $\mathbf{H}$ are block-diagonal matrices with one block per study, and each of the terms $\boldsymbol{V}_{G_0,G}^\top \mathbf{V}_{G_0}^{-1} \boldsymbol{U}_{G_0}$, $\boldsymbol{V}_{G_0,G}^\top \mathbf{V}_{G_0}^{-1} \boldsymbol{V}_{G_0,G}$, and $\boldsymbol{U}_{G_0}^\top \mathbf{V}_{G_0}^{-1} \boldsymbol{U}_{G_0}$ can be written as a sum over terms calculated separately within each study.

**3. If $\epsilon \sim \mathcal{N}(\mathbf{0}, \mathbf{\Omega})$, then 1. and 2. apply after the following transformation:** Premultiply $\mathbf{\Omega}^{-1/2}$ to each of $\boldsymbol{Y}$, $\mathbf{C}$, $\mathbf{G}_0$, and $\boldsymbol{G}$ to obtain generalized least squares (GLS) slopes and standard errors for $\boldsymbol{G}$. This applies when $\mathbf{\Omega}$ is a known matrix, but in practice is often used with $\hat{\mathbf{\Omega}}$ estimated under the null hypothesis (genetic effect $\beta = 0$, and technical covariate effects unconstrained) in genome-wide association studies (GWAS), and the effect of an individual variant will be small in magnitude. This approach is used in APEX for multiple-variant xQTL analysis.

#### 0.7 xQTL Summary Data Storage Formats

APEX generates compact storage files for score statistics $\mathbf{U}_s = \mathbf{G}_s^\top \mathbf{\Omega}_s^{-1}(\mathbf{I} - \mathbf{H}_s)\mathbf{Y}_s$ and $\mathbf{V}_s = \mathbf{G}_s^\top \mathbf{\Omega}_s^{-1}(\mathbf{I} - \mathbf{H}_s)\mathbf{G}_s$ for data sharing and meta-analysis while maintaining genetic privacy. For *cis* analysis, we store elements of $\mathbf{U}_s$ for variants within a window (1 Mbp upstream or downstream by default) of each molecular trait in indexed and compressed tab-delimited text files. For $\mathbf{V}_s$, we store all variant pairs within a specified sliding window across the genome, taken as twice the size of the specified window size for cis-xQTL analysis. We decompose $\mathbf{V} = \mathbf{G}^\top \mathbf{A} \mathbf{G} + \mathbf{G}^\top \mathbf{B} \mathbf{B}^\top \mathbf{G}$, where $\mathbf{A}$ is a sparse diagonal matrix and $\mathbf{B}^\top \mathbf{G}$ is a dense matrix of size $m \times q$. $\mathbf{B}^\top \mathbf{G}$ is stored in a BGZIP block-compressed and indexed text file together with variant summary data. $\mathbf{G}^\top \mathbf{A} \mathbf{G}$ is stored separately using a fixed-width binary integer format, which is block-compressed using XZ format and indexed by byte offset for efficient random access by region. Further details are available online.

### References

Aguet, François et al. (2019). "The GTEx Consortium atlas of genetic regulatory effects across human tissues". In: *BioRxiv*, p. 787903.

Brent, Richard P (1973). "Algorithms for minimization without derivatives". In: ISSN: 0486419983.

Chen, Han and Matthew P Conomos (2020). "GMMAT-package: Generalized Linear Mixed Model Association Tests". In:

Chen, Han et al. (2019). "Efficient variant set mixed model association tests for continuous and binary traits in large-scale whole-genome sequencing studies". In: *The American Journal of Human Genetics* 104.2, pp. 260–274. ISSN: 0002-9297.

Consortium, GTEx (2017). "Genetic effects on gene expression across human tissues". In: *Nature* 550.7675, pp. 204–213. ISSN: 1476-4687.

Feng, Shuang et al. (2014). "RAREMETAL: fast and powerful meta-analysis for rare variants". In: *Bioinformatics* 30.19, pp. 2828–2829. ISSN: 1460-2059.

Giesbrecht, Francis G and Joseph C Burns (1985). "Two-stage analysis based on a mixed model: large-sample asymptotic theory and small-sample simulation results". In: *Biometrics*, pp. 477–486.

Gogarten, Stephanie M et al. (2019). "Genetic association testing using the GENESIS R/Bioconductor package". In: *Bioinformatics* 35.24, pp. 5346–5348.

Halekoh, Ulrich, Søren Højsgaard, et al. (2014). "A kenward-roger approximation and parametric bootstrap methods for tests in linear mixed models–the R package pbkrtest". In: *Journal of Statistical Software* 59.9, pp. 1–30.

Harville, David A (1974). "Bayesian inference for variance components using only error contrasts". In: *Biometrika* 61.2, pp. 383–385.

Henderson, Charles R (1973). "Sire evaluation and genetic trends". In: *Journal of Animal Science* 1973.Symposium, pp. 10–41.

Hrong-Tai Fai, Alex and Paul L Cornelius (1996). "Approximate F-tests of multiple degree of freedom hypotheses in generalized least squares analyses of unbalanced split-plot experiments". In: *Journal of statistical computation and simulation* 54.4, pp. 363–378.

Jiang, Longda et al. (2019). *A resource-efficient tool for mixed model association analysis of large-scale data.* Tech. rep. Nature Publishing Group.

Kang, Hyun Min et al. (2010). "Variance component model to account for sample structure in genome-wide association studies". In: *Nature genetics* 42.4, pp. 348–354. ISSN: 1546-1718.

Kenward, Michael G and James H Roger (1997). "Small sample inference for fixed effects from restricted maximum likelihood". In: *Biometrics*, pp. 983–997. ISSN: 0006-341X.

Kuznetsova, Alexandra, Per B Brockhoff, and Rune HB Christensen (2017). "lmerTest package: tests in linear mixed effects models". In: *Journal of statistical software* 82.13, pp. 1–26. ISSN: 1548-7660.

Kuznetsova, Alexandra, Per B Brockhoff, Rune HB Christensen, et al. (2017). "lmerTest package: tests in linear mixed effects models". In: *Journal of statistical software* 82.13, pp. 1–26.

Lawley, Derrick N (1940). "Vi.âthe estimation of factor loadings by the method of maximum likelihood". In: *Proceedings of the Royal Society of Edinburgh* 60.1, pp. 64–82. ISSN: 0370-1646.

Lippert, Christoph et al. (2011). "FaST linear mixed models for genome-wide association studies". In: *Nature methods* 8.10, pp. 833–835. ISSN: 1548-7105.

Liu, Yaowu and Jun Xie (2020). "Cauchy combination test: a powerful test with analytic p-value calculation under arbitrary dependency structures". In: *Journal of the American Statistical Association* 115.529, pp. 393–402. ISSN: 0162-1459.

Loh, Po-Ru et al. (2015). "Efficient Bayesian mixed-model analysis increases association power in large cohorts". In: *Nature genetics* 47.3, p. 284. ISSN: 1546-1718.

Owen, Art B and Jingshu Wang (2016). "Bi-cross-validation for factor analysis". In: *Statistical Science* 31.1, pp. 119–139. ISSN: 0883-4237.

Patterson, H Desmond and Robin Thompson (1971). "Recovery of inter-block information when block sizes are unequal". In: *Biometrika* 58.3, pp. 545–554.

Price, Alkes L et al. (2006). "Principal components analysis corrects for stratification in genome-wide association studies". In: *Nature genetics* 38.8, pp. 904–909. ISSN: 1546-1718.

Satterthwaite, Franklin E (1946). "An approximate distribution of estimates of variance components". In: *Biometrics bulletin* 2.6, pp. 110–114. ISSN: 0099-4987.

Stegle, Oliver et al. (2012). "Using probabilistic estimation of expression residuals (PEER) to obtain increased power and interpretability of gene expression analyses". In: *Nature protocols* 7.3, p. 500. ISSN: 1750-2799.

Svishcheva, Gulnara R et al. (2012). "Rapid variance componentsâbased method for whole-genome association analysis". In: *Nature genetics* 44.10, pp. 1166–1170. ISSN: 1546-1718.

Wang, Jingshu (2016). "FACTOR ANALYSIS FOR HIGH-DIMENSIONAL DATA". Thesis.

Welch, Bernard L (1947). "The generalization ofstudent's' problem when several different population variances are involved". In: *Biometrika* 34.1/2, pp. 28–35. ISSN: 0006-3444.

Zhan, Xiaowei et al. (2016). "RVTESTS: an efficient and comprehensive tool for rare variant association analysis using sequence data". In: *Bioinformatics* 32.9, pp. 1423–1426. ISSN: 1460-2059.

Zhou, Wei et al. (2018). "Efficiently controlling for case-control imbalance and sample relatedness in large-scale genetic association studies". In: *Nature genetics* 50.9, pp. 1335–1341. ISSN: 1546-1718.

Zhou, Xiang and Matthew Stephens (2012). "Genome-wide efficient mixed-model analysis for association studies". In: *Nature genetics* 44.7, pp. 821–824. ISSN: 1546-1718.
