## Supplementary Figures for "A versatile toolkit for molecular QTL mapping and meta-analysis at scale"

### Supplementary Figure 1. Correlation structure of simulated expression across genes

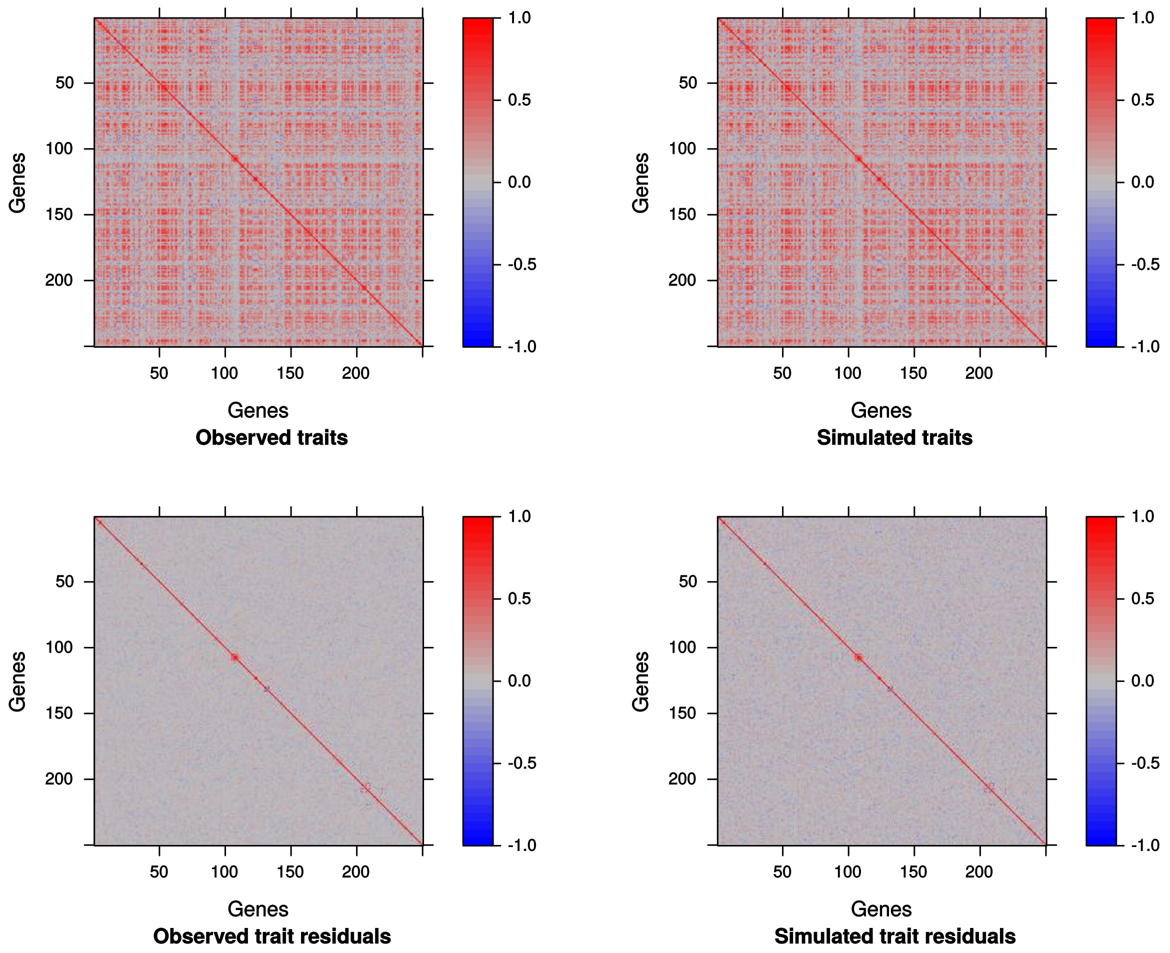


We simulated expression data in the Geuvadis cohort with realistic correlation structures. Here, we show the covariance matrices of observed (left) and simulated (right) expression data across genes before and after regressing out technical covariates (top vs bottom), which include genotype PCs and expression PCs (ePCs). ePCs are re-calculated for each simulated data set; here we show a single simulation replicate as an example. We use the covariance between observed expression and technical covariates (other than ePCs) and the complete singular value decomposition of observed expression to simulate gene expression under a multivariate normal distribution. Simulated data closely match the correlation structure of observed expression across genes.

### Supplementary Figure 2. Correlation structure of simulated expression across individuals

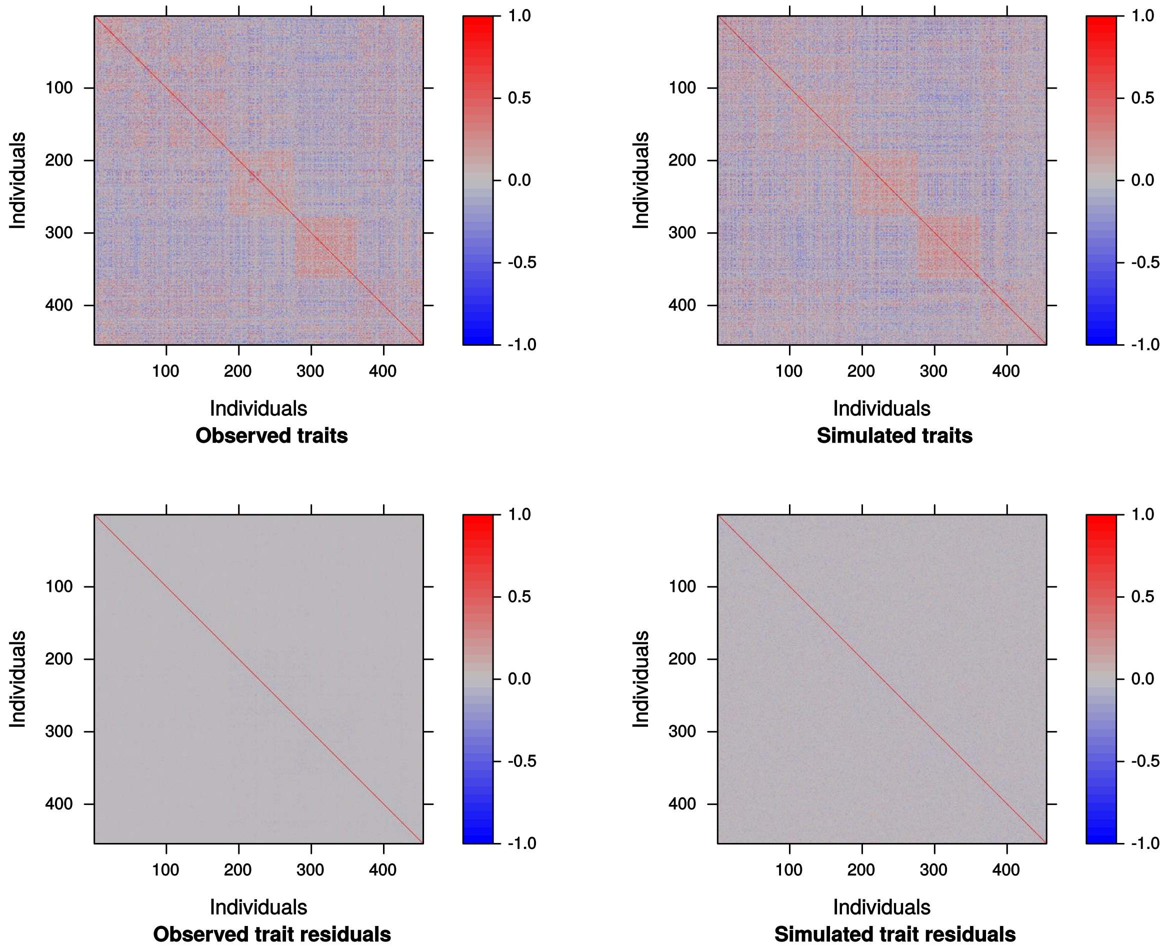


We simulated expression data in the Geuvadis cohort with realistic correlation structures. Here, we show the covariance matrices of observed (left) and simulated (right) expression data across individuals before and after regressing out technical covariates (top vs bottom), which include genotype PCs and expression PCs (ePCs). ePCs are re-calculated for each simulated data set; here we show a single simulation replicate as an example. We use the covariance between observed expression and technical covariates (other than ePCs) and the complete singular value decomposition of observed expression to simulate gene expression under a multivariate normal distribution. Simulated data closely match the correlation structure of observed expression across individuals.

**Supplementary Figure 3. Empirical Type I error rates for single-variant xQTL association tests with factor covariates modeled as fixed or random effects**

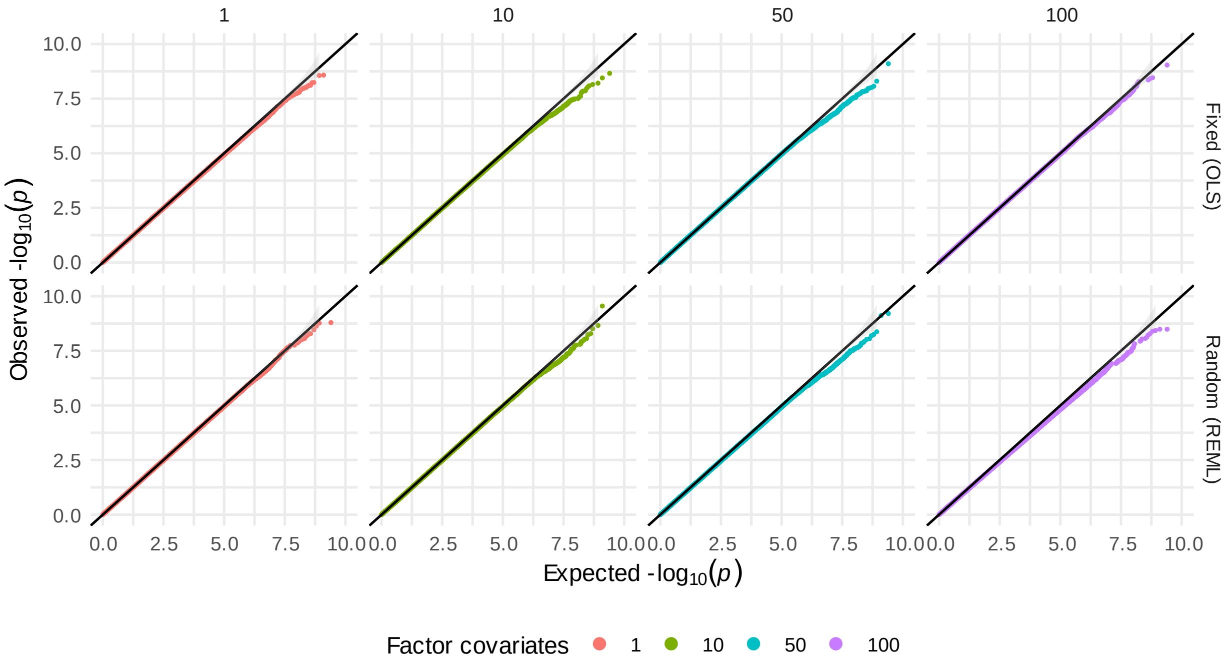


We simulated expression traits under the null hypothesis in the Geuvadis cohort (n = 454 participants). In each of 100 replicates, 17,815 expression traits were simulated using the covariance between observed expression and covariates (sex, genotype PCs, and self-reported ancestry group), and the residual covariance of observed expression traits as described in Methods. In each replicate, we re-calculated common factors (eFA or ePC) from the simulated expression matrix. Shown are Q-Q plots of single-variant association p-values across replicates when 1, 10, 50, or 100 factor covariates are included as fixed (upper) or random (lower) effects.

### Supplementary Figure 4. Efficient linear mixed models with tens of thousands of traits

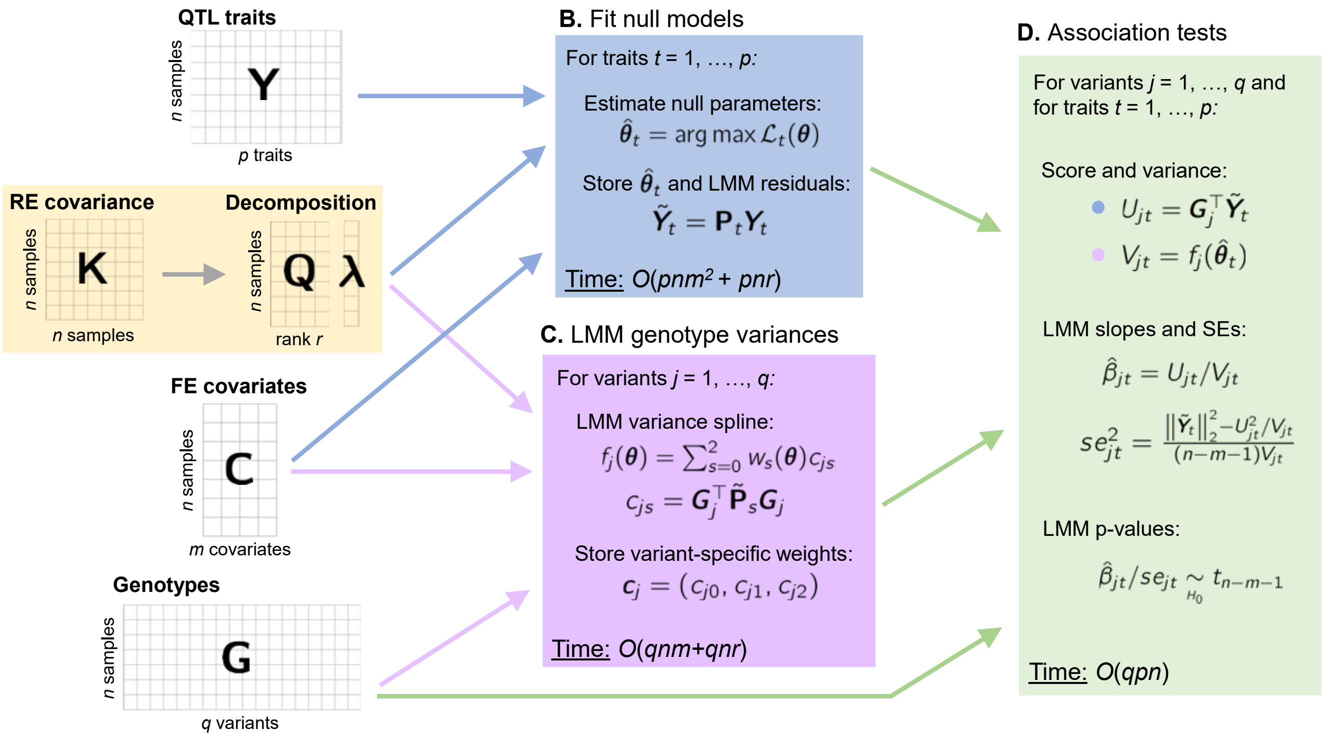


APEX uses a three-stage approach to efficiently perform linear mixed model (LMM) association analysis with tens of thousands of traits. Here, we show a low-rank covariance matrix; APEX also supports full-rank sparse, block diagonal, and dense covariance matrices (e.g., genetic relatedness matrices; GRMs). Given the decomposed matrix, we estimate and store variance component parameters and LMM trait residuals under the null hypothesis (no single-variant effects) for each trait (Panel B). Next, we calculate and store spline interpolations terms for the residual variances of genotypes (Supplementary Materials) as a function of variance component parameters (Panel C). Finally, we use these data to calculate LMM association tests in linear time (Panel D).

### Supplementary Figure 5. LMM software concordance

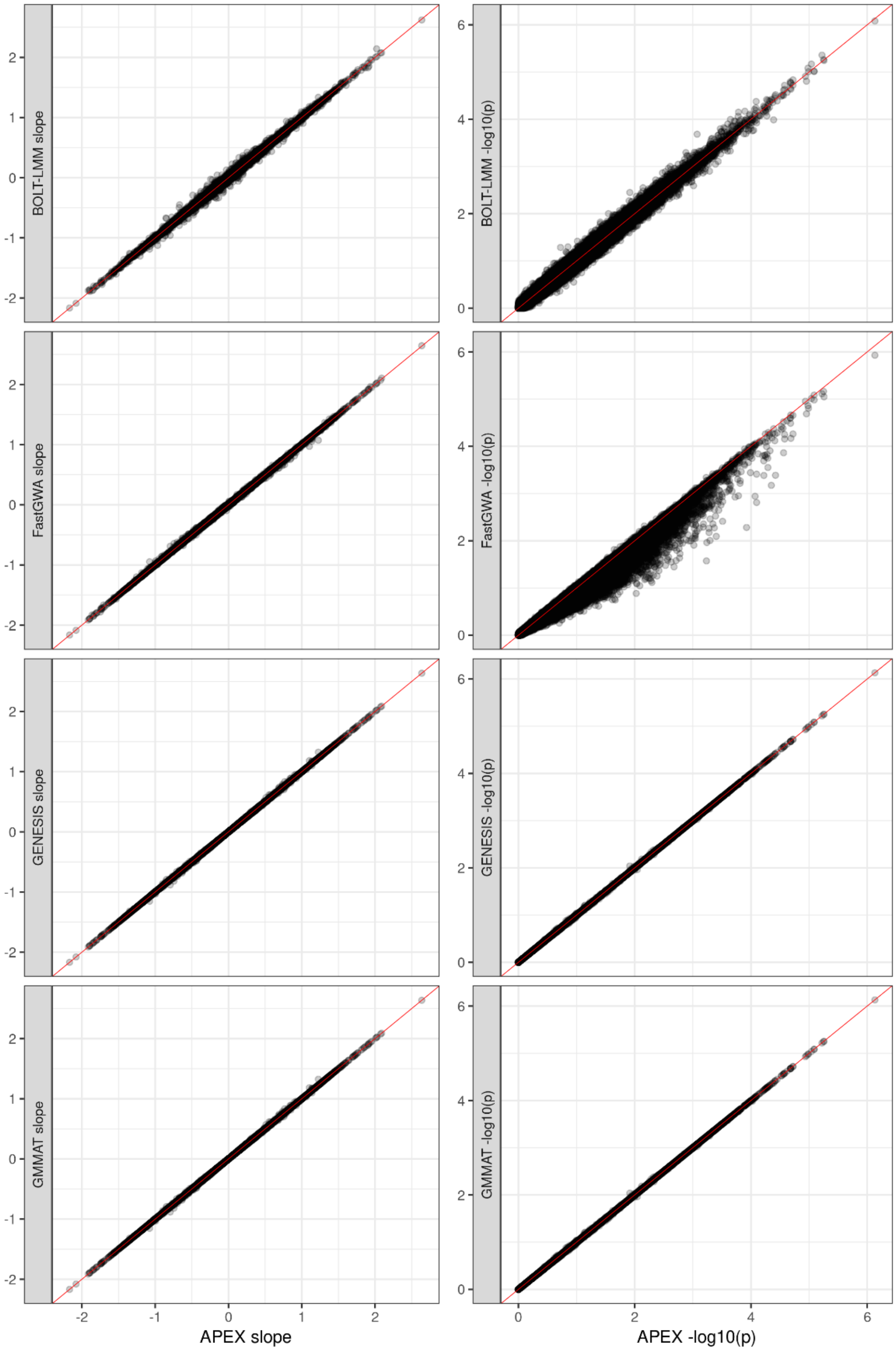


Concordance of slope estimates and p-values from linear mixed models (LMMs) across software. Here, we used genotype and covariates data for a subset of 10,000 participants from the UK Biobank and simulated molecular trait data (Methods) to compare computational requirements for xQTL LMM association analysis using APEX, GMMAT, GENESIS, BOLT-LMM, and FastGWAS. LMM association tests from FastGWAS and BOLT-LMM use approximations that rely on large sample sizes; APEX, GMMAT, and GENESIS use no such approximations. Shown are LMM single-variant slope estimates (left) and p-values (right) for each tool across 100 simulated traits and variants on chromosome 22.

**Supplementary Figure 6. Concordance of ACAT and permutation-based eGene p-values**

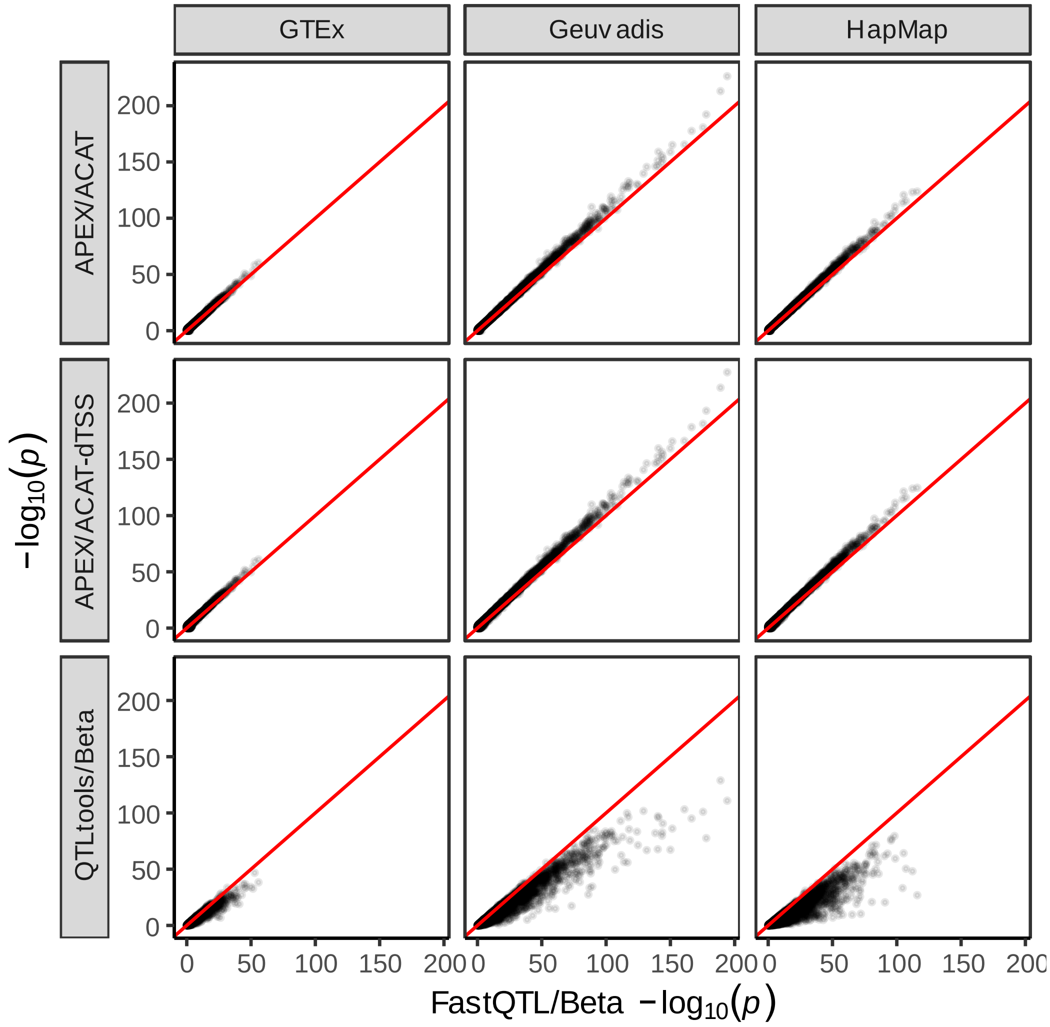


Concordance of eGene p-values in each LCL eQTL study calculated using unweighted or dTSS-weighted ACAT (implemented in APEX), beta-approximated permutation omnibus p-values implemented in QTLtools (Delaneau et al., 2017) and FastQTL (Ongen et al., 2016). QTLtools used simple linear regression with expression residuals (SLR-resid), whereas other tools use multiple linear regression (MLR) to adjust for technical covariates. As a consequence, QTLtools p-values appear deflated relative to other tools. We note that QTLtools can also perform MLR by constructing a VCF with genotype residuals.

Here (using empirical expression), ACAT p-values are generally slightly more significant than FastQTL beta p-values, and both are generally more significant than QTLtools SLR-resid beta p-values. Similarly, in simulation studies under the null hypothesis (shown in Main Figure 3, panel B), ACAT p-values were well-calibrated, while FastQTL beta p-values were slightly conservative, and QTLtools SLR-resid beta p-values were severely conservative.

**Supplementary Figure 7. Accurate and flexible multiple-variant QTL meta-analysis**

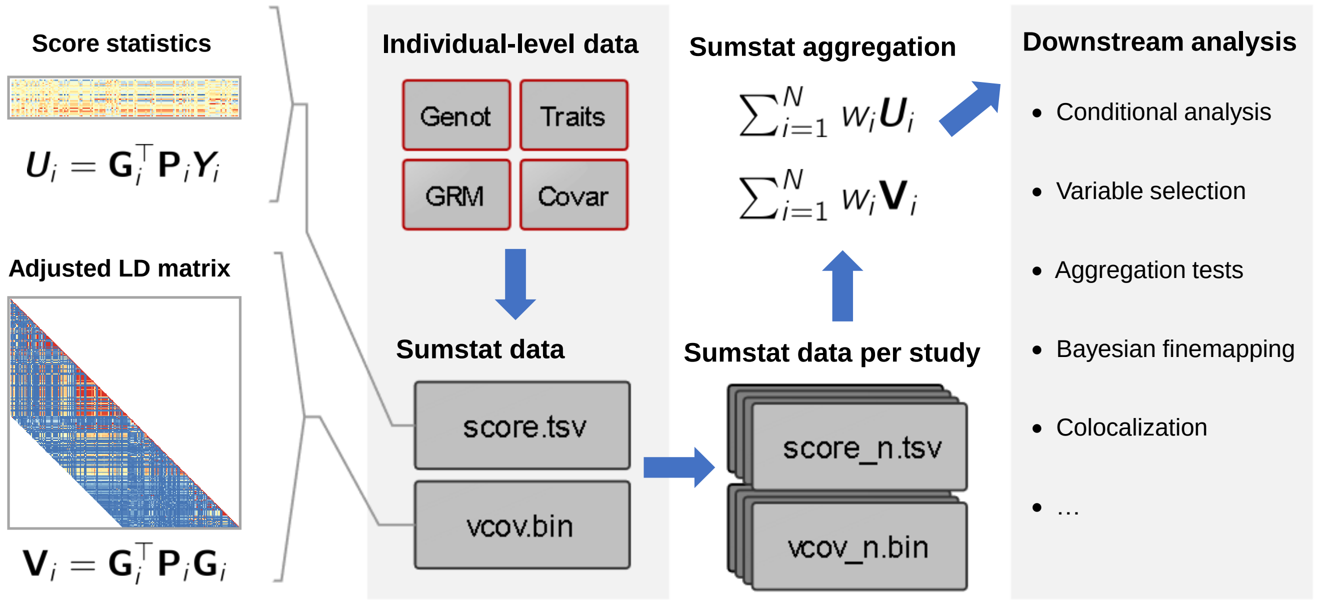


APEX enables data-sharing and meta-analysis via compact summary data files. We provide condensed association summary statistic ("score") and variance-covariance data ("vcov") files, which enable a wide range of joint multiple-variant meta-analysis procedures. Joint/conditional and stepwise regression procedures to identify secondary QTL signals and multiple-variant aggregation tests to from summary data are directly implemented in APEX. Score and vcov files generated by APEX further enable Bayesian finemapping, colocalization analysis, penalized regression, and other analyses from QTL meta-analysis summary statistics using external software.

### Supplementary Figure 8. APEX vcov storage format

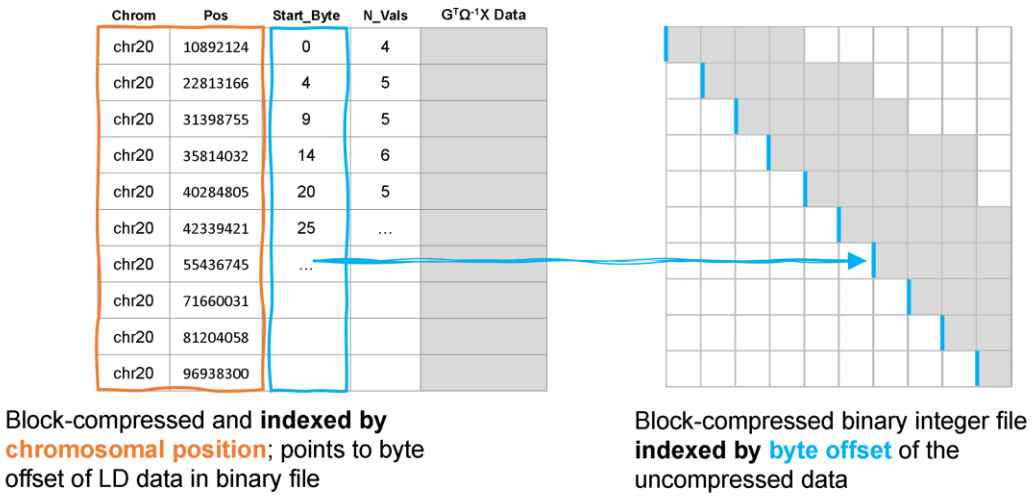


Visualization of QTL variance-covariance data format implemented in APEX. In one file, we store the product of genotypes and technical covariates (left), which is block-compressed indexed by genomic position. Separately, we store the cross product matrix of genotypes (right), which is stored as a symmetric band diagonal matrix in a block-compressed binary file indexed by byte offset. For the cross product matrix, we use a 2 Mbp window by default, or twice the specified *cis*-eQTL window size (1 Mbp by default). This indexing strategy enables rapid querying by region or variant.

### Supplementary Figure 9. Secondary signal discovery flowchart

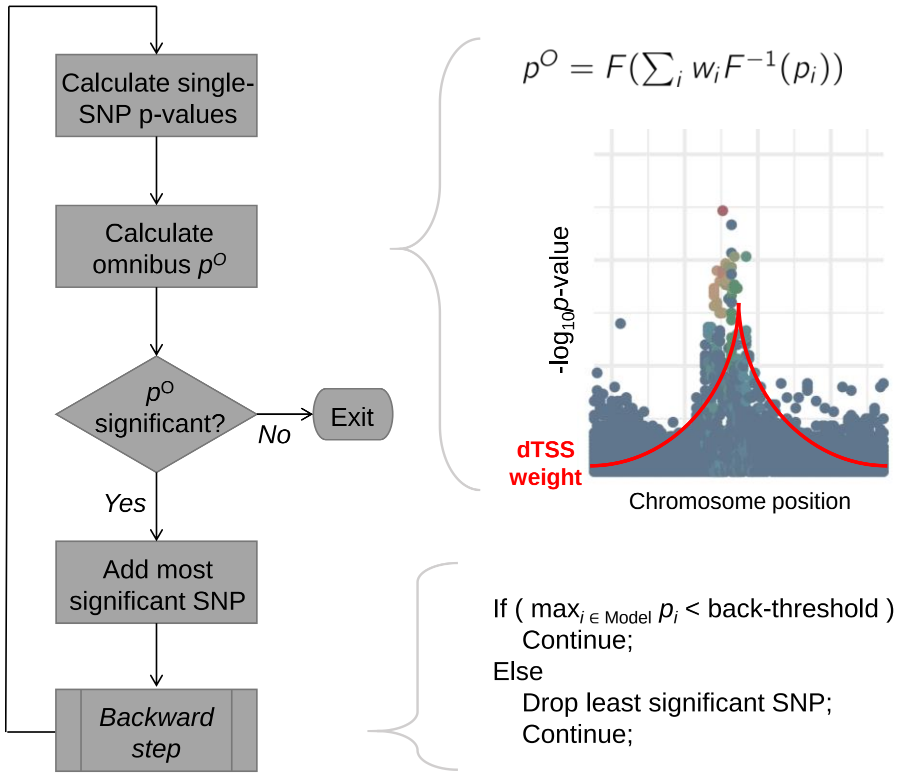


Secondary signal detection algorithms in APEX. At each iteration, we calculate an omnibus p-value testing the null hypothesis that the additive effects of all remaining variant are 0 conditional on variants already included in the model. If the omnibus test is significant, the most significant remaining variant is added to the model. Optionally, we then check joint p-values of all variants in the model, and drop the least significant variant from the model if its joint p-value falls below a threshold. Further details are given in Supplementary Materials.

**Supplementary Figure 10. QTL Studies Conducted by Year**

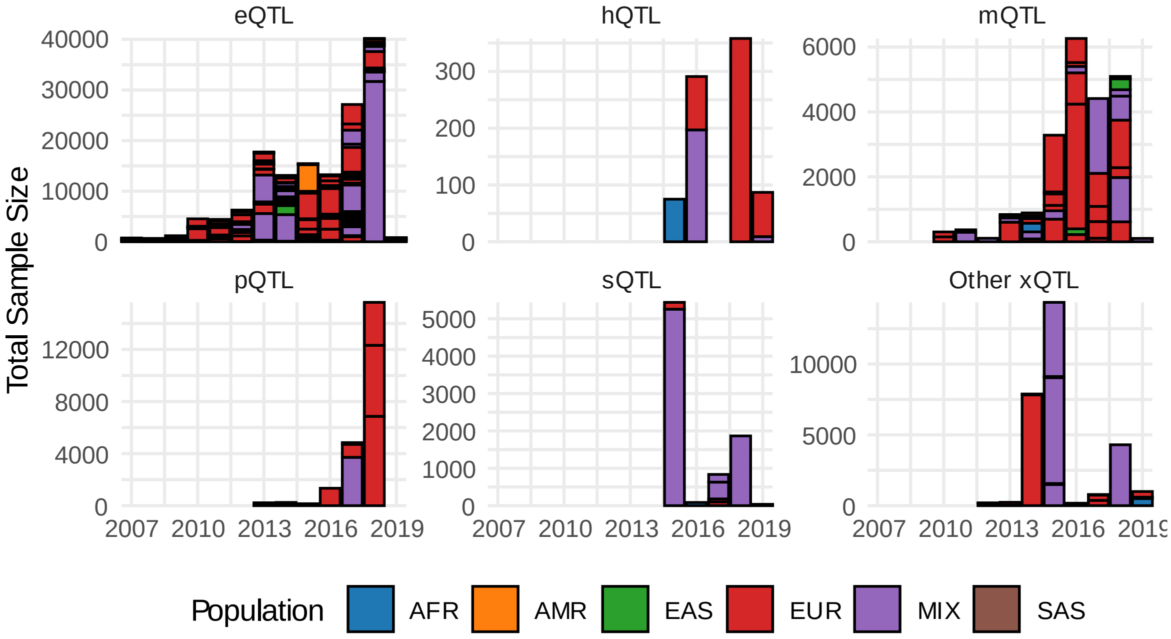


Sample sizes of expression QTL (eQTL), histone QTL (hQTL), protein QTL (pQTL), splicing QTL (sQTL), and other molecular QTL (QTL) association studies by year. Individual studies are delineated by black outline, and colored by study population. Data obtained from QTLbase (Zheng et al., 2020).

The vast majority of QTL study data are not publicly available, which hampers mega- or meta-analysis. LD reference panels are not well-suited for summary-based analysis of studies that include multiple and/or understudied ethnicities, or studies with small sample sizes. A large fraction of QTL studies feature multiple and/or non-European ethnic groups, and most have <1000 samples.

**Supplementary Figure 11. Enrichment of LCL eQTLs in tissue-specific DNase I hypersensitive sites**

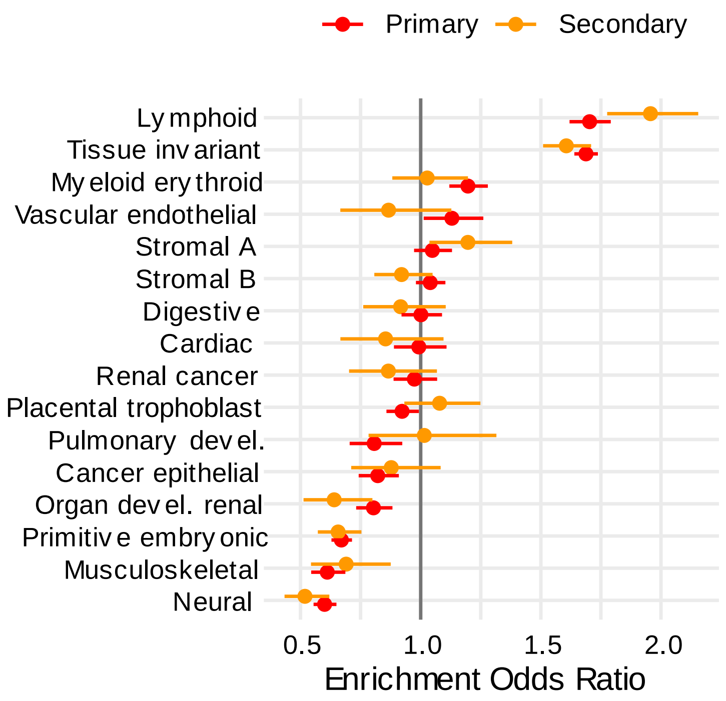


Primary and secondary LCL eQTL enrichment in tissue-specific DNase I hypersensitive sites (DHSs) (Meuleman et al., 2020), adjusted for minor allele frequency (MAF) (log-transformed cubic b-spline) and distance to nearest transcription start site (TSS) (log-transformed cubic b-spline) as described in Methods. Shown are enrichment odds ratios (exponentiated logistic regression coefficients, ± 2 standard errors) estimated separately by including a single DHS tissue category per model.

**Supplementary Figure 12. Enrichment of LCL eQTLs in NHGRI-EBI GWAS Catalog**

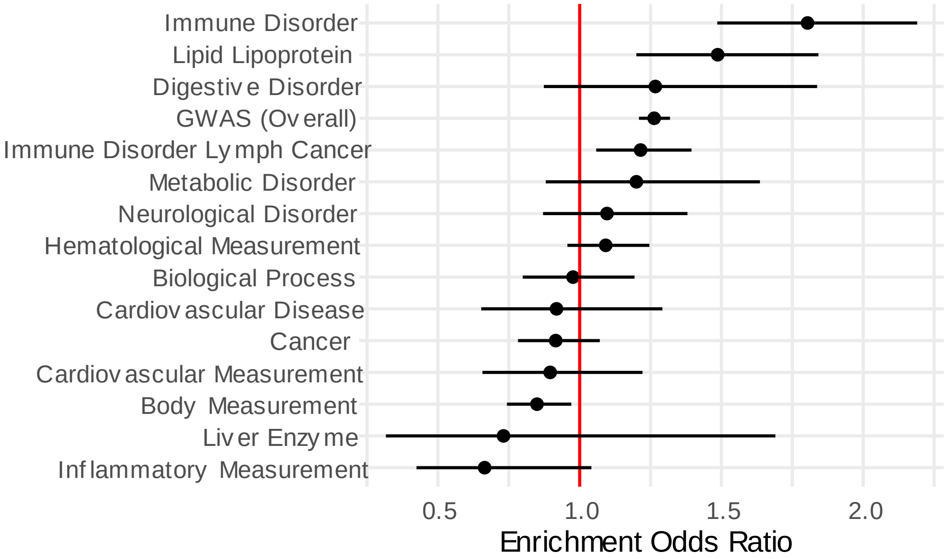


LCL eQTL enrichment for categories of traits in the NHGRI-EBI GWAS Catalog, adjusted for minor allele frequency (MAF) (log-transformed cubic b-spline) and distance to nearest transcription start site (TSS) (log-transformed cubic b-spline) as described in Methods. LCL eQTLs show strongest enrichment with immune disorders. Shown are enrichment odds ratios (exponentiated logistic regression coefficients, ± 2 standard errors) estimated separately by including a single GWAS trait category per model.
