## Supplementary Tables for "A versatile toolkit for molecular QTL mapping and meta-analysis at scale"

**Supplementary Table 1. Benchmarking LMM association analysis of 16K traits across 10K samples**

| | CPU hours | Max memory |
|---|---|---|
| APEX (p-value < 1e-5) | 7.5 | 4.88 GB |
| APEX | 20.8 | 4.88 GB |
| FastGWA | 52.1 | 0.14 GB |
| BOLT-LMM | 1,068.9 | 0.67 GB |
| GMMAT (approx.) | 8,975.6 | 0.77 GB |
| GENESIS (approx.) | 8,492.7 | 6.86 GB |

Time and memory for genome-wide LMM association analysis of 16,329 simulated molecular traits across 590,606 SNPs using genotype data from a sample of 10,000 participants from the UK Biobank, controlling for 10 covariates. All reported times include loading data, estimating LMM null models, calculating and writing singe-variant association tests. All jobs were run using a single CPU core.

For "APEX (p-value < 1e-5)", all SNP-trait associations are calculated internally, and only p-value < 1e-5 associations are recorded in output files (using –pvalue command line option). For "APEX" and all subsequent rows, all SNP-trait associations are recorded in output files.

For GMMAT and GENESIS, we analyzed a subset of 100 simulated traits, and multiplied the total CPU hours for 100 traits by a factor of 163.29 to estimate CPU hours for all 16,329 traits. For all other methods (APEX, FastGWA, and BOLT-LMM), we analyzed all 16,329 traits.

**Supplementary Table 2. APEX sumstat and vcov benchmarking**

| | GTEx v8 LCLs | Geuvadis | HapMap |
|---|---|---|---|
| **Storage size** | | | |
| vcov file size | 20.88 Gb | 32.62 Gb | 116.84 Gb |
| vcov index file size | 0.60 Gb | 1.72 Gb | 2.89 Gb |
| cis sumstat file size | 0.49 Gb | 0.42 Gb | 0.47 Gb |
| **Time and memory** | | | |
| vcov files: CPU time | 32.08 hrs | 62.04 hrs | 75.30 hrs |
| vcov files: memory usage | 0.59 Gb | 1.24 Gb | 1.58 Gb |
| cis sumstat files: CPU time | 0.17 hrs | 0.27 hrs | 0.33 hrs |
| cis sumstat files: memory usage | 0.81 Gb | 2.14 Gb | 2.43 Gb |
| **Study characteristics** | | | |
| Variants | 10,932,660 | 10,945,700 | 10,943,352 |
| Genotypes | Hard-calls (GT) | Hard-calls (GT) | Dosages (DS) |
| Sample size | 147 | 454 | 518 |
| Covariates | 23 | 70 | 97 |
| Genes | 22,759 | 17,815 | 16,329 |

Storage size, CPU time, and memory usage to generate APEX sumstat and vcov files across all autosomes with 2 Mbp sliding window size in each study.  All jobs were run using a single CPU.

**Supplementary Table 3. APEX vcov vs RareMetalWorker cov files for chromosome 20**

| | GTEx v8 LCLs | Geuvadis | HapMap |
|---|---|---|---|
| **Storage size** | | | |
| APEX vcov + index files | 0.508 GB | 0.808 GB | 2.771 GB |
| RareMetalWorker cov file (gzip) | 10.910 GB | 12.390 GB | 12.570 GB |
| **CPU time** | | | |
| APEX vcov + index | 0.734 hrs | 1.049 hrs | 1.437 hrs |
| RareMetalWorker | 1.597 hrs | 2.114 hrs | 2.197 hrs |
| **Memory usage** | | | |
| APEX vcov + index | 0.237 GB | 0.406 GB | 0.501 GB |
| RareMetalWorker | 0.136 GB | 0.163 GB | 0.169 GB |

Comparison of storage size, CPU time, and memory usage to generate adjusted LD in 2 Mbp sliding windows for chromosome 20 using APEX and RareMetalWorker (RMW) version 4.15.1. All jobs were run using a single CPU. RMW stores adjusted LD using tab- and comma-delimited text files; APEX stores LD using tab-delimited index files and binary integer vcov files. RMW files were gzip compressed ( RMW “--zip" option). APEX vcov files are internally xz block-compressed and indexed.
